# Estimating maximum sustainable yield of snow crab (*Chionoecetes opilio*) off Tohoku Japan via a state-space assessment model with time-varying natural mortality

**DOI:** 10.1101/2020.02.02.931428

**Authors:** Yasutoki Shibata, Jiro Nagao, Yoji Narimatsu, Eisuke Morikawa, Yuto Suzuki, Shun Tokioka, Manabu Yamada, Shigeho Kakehi, Hiroshi Okamura

## Abstract

Yield from fisheries is a tangible benefit of ecosystem services and sustaining or restoring a fish stock level to achieve a maximum sustainable yield (MSY). Snow crab (*Chionoecetes opilio*) off Tohoku has been managed by a total allowable catch since 1996, although their abundance has not increased even after 2011, when fishing pressure rapidly decreased because of the Great East Japan Earthquake. This implies that their biological characteristics, such as recruits, natural mortality coefficient (*M*), and terminal molting probabilities (*p*), might have changed. We developed “just another state-space stock assessment model (JASAM)” to estimate the MSY of the snow crab off Tohoku, Japan, considering interannual variations in *M* and *p*. The multi-model inference revealed that *M* increased from 0.2 in 1997 to 0.59 in 2018, although it was not different among the instars, sex, nor terminal molt of crabs. The parameter *p* also increased by 1.34–2.46 times depending on the instar growth stages from 1997 to 2018. We estimated the MSYs in three scenarios, which drastically changed if *M* and *p* were set as they were in the past or at the current values estimated from this study. This result indicated that the MSY of snow crab would also be time-varying based on their time-varying biological characteristics.

## Introduction

Ecosystem services are the benefits that nature can provide to households, communities, and economies (Boyd and Banzhaf 2006), and fish production (yield) is a tangible benefit of ecosystem services (Tomscha and Gergel 2016). Although the development of fishery and transportation technologies has become a factor to increase yield, an increase in yield may lead to overfishing and reduced yield. At the United Nations summit in 2015, the Sustainable Development Goals (SDGs) was adopted in “The 2030 Agenda for Sustainable Development” as a common international objective from 2016 to 2030 (UN General Assembly 2015a). The SDGs comprise 17 goals and 169 targets, and maintaining or restoring fish stock levels to achieve a maximum sustainable yield (MSY) as determined by their biological characteristics is one of main targets of the SDGs (UN General Assembly 2015b).

In Japan, the Fisheries Law has been amended for the first time in 70 years since it was enacted in 1949. In the amended Fisheries Law, it is necessary to set a target for fish abundance that maintains the MSY of a (spawning stock) biomass calculated under natural conditions in the present and reasonably foreseeable future (Fisheries Agency; https://www.jfa.maff.go.jp/j/kikaku/kaikaku/attach/pdf/suisankaikaku-20.pdf last accessed 29 January 2020). Some fish stocks that are legally managed by regulated total allowable catch (TAC) have begun to be managed under this amended Fisheries Law.

Snow crab (*Chionoecetes opilio*) off the Tohoku region (Figure 1) have been managed by a total allowable catch (TAC) since 1996, and are one of the most valuable exploited species in Japan. In snow crab, terminal molting stops body growth as it matures (Yoshida 1941; Conan and Comeau 1986). At present, it is assumed that snow crab off Tohoku molt once a year in and after instar VI, and then molt finally from instar X; this is based on a case study in which specimens from the Sea of Japan were reared (Kuwahara et al. 1995).

**Fig. 1.**
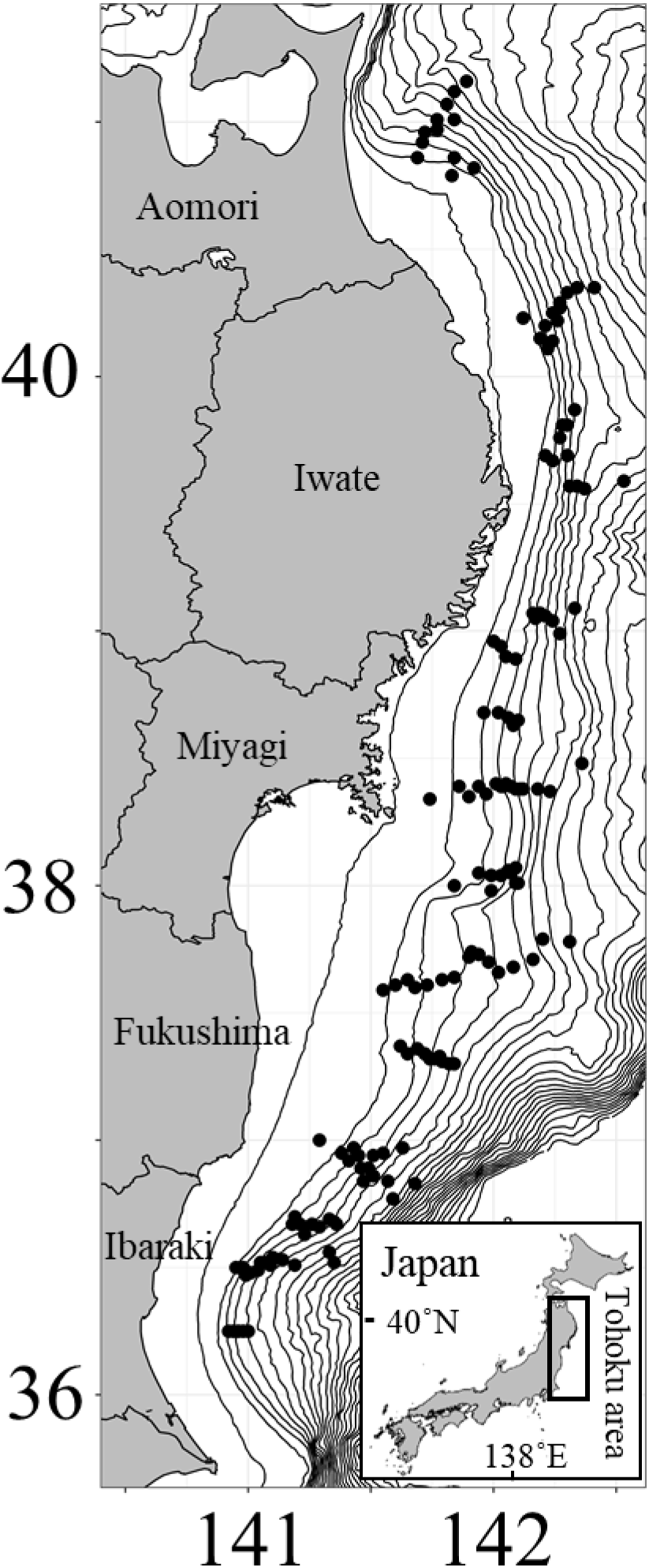
Stations surveyed by the R/V Wakataka-maru since 1997 (black circles, 150 stations). The survey area covered off Aomori, Iwate, Miyagi, Fukushima, and Ibaraki prefectures from 150 to 900 m depth. Contours drawn by 100 m depth.

Although total landings in the Tohoku area were around 100 gross ton before the 2011, total landings and fishing efforts (the number of tows that caught at least one snow crab by bottom trawl vessels) rapidly decreased in 2012 (Figure 2) because of the Great East Japan Earthquake and tsunami in March 2011, which destroyed much of the fisheries-related infrastructure, such as fishing vessels, fishing ports, and marine product processing factories. Fishing efforts for snow crab after the earthquake have been less than 2% of those before 2011 (Shibata et al. 2019). Regarding bottom trawl fishing off Fukushima, only trial fishing has been carried out since 2011 (Shibata et al. 2017).

**Fig. 2.**
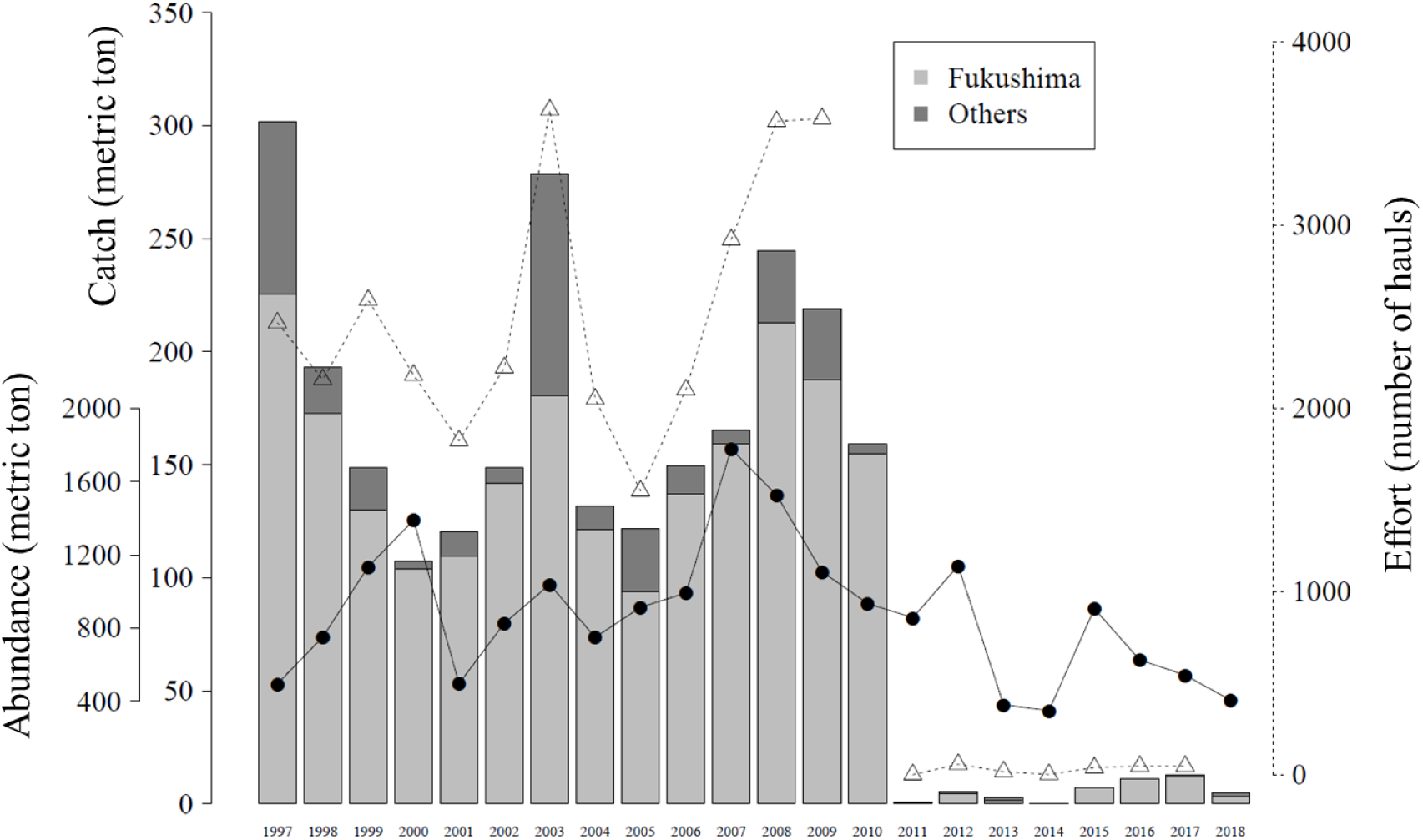
Total catch (metric ton, bar graph), estimated abundance (metric ton) by a swept area method from survey data of the R/V Wakataka-maru (black circles) and effort of bottom trawl vessels (number of hauls, white triangles). Total catch was distinguished between that of Fukushima (gray) and other prefectures (dark gray). Effort value in 2018 was under calculation.

The stock status of snow crab has been assessed by scientists of the Japanese Fisheries Research and Education Agency (FRA), based on their estimated abundance (fishable biomass, males: only for carapace width (*cw*) ≥ 80 mm, females: only for matured) by a swept area method from the survey results of a research vessel (R/V) every year. Regardless of the quite low fishing efforts, the observed abundance has continued decreasing rather than increasing, contrary to expectations (Figure 2). Because fishing pressures have been kept quite low since March 2011, other factors have caused this continued decrease. In fact, the two-year projected abundances that were needed to calculate the allowable biological catch (ABCs) had likely been overestimated since 2012 by a previously developed model (Ueda et al. 2009), with the fishing mortality coefficient (*F*) set as almost zero (Shibata et al. 2019). One possible reason for this overestimation is that the biological characteristics, such as recruits, natural mortality coefficient (*M*), and terminal molting probability (*p*), could have changed in recent years. Especially, *M* and *p* might have increased and maintained high values; the previous model assumed that *M* and *p* did not vary with time.

Since we are interested in the time variation of the parameters, it is necessary to estimate these parameters using an appropriate statistical model. In previous studies, these parameters were treated as constant with time, or expressed with random effects (Yamasaki 1988; Szuwalski and Turnock 2016; Murphy et al. 2018). However, it is more natural to assume that the parameters change continuously rather than randomly changing every year. Therefore, we assumed that the biological characteristic parameters followed a random walk (RW). A state-space stock assessment model (SAM) has been reported (Nielsen and Berg 2014), in which some parameters vary by an RW process. Although an age-structured model, such as statistical catch at age (SCAA), usually assumes that a selectivity pattern is constant over time (Butterworth and Rademeyer 2008), SAM can estimate a time-varying selectivity following an RW with a multivariate normal distribution (Nielsen and Berg 2014). SAM also enables the use of information criteria, such as Akaike’s information criterion (AIC) (Akaike 1974), for model selection, as opposed to the penalized likelihood approach (Nielsen and Berg 2014). Since the model developed in this study has some structures in common with SAM, it will be called JASAM (just another SAM).

The purpose of this study is to develop a state-space model that considers variations and predicts the future abundance based on the stock–recruitment relationship defined in this study, thereby estimating the MSY of snow crab off the Tohoku region, Japan, taking into account interannual variations in biological characteristic parameters, such as recruits, *M,* and *p*.

## Materials and Methods

### Scientific bottom trawl surveys

To estimate the abundance of snow crab, scientific surveys using a bottom trawl net by the R/V Wakataka-maru have been carried out between 1997 and 2018 on the northern part of Honshu Island, Japan (Tohoku region [Figure 1]). A total of 150 survey stations were set to tow at depths from 150 to 900 m from September to November, where the spatial distribution of snow crab was at depths ranging from 150 to 700 m and the main distribution range of fishable crabs (males with *cw* ≥ 80 mm and mature females) is approximately 400–550 m (Kitagawa 2000). The total length and mouth width of the trawl net were 44.1 m and 5.4 m, respectively. The mesh size of the net was 50 mm and a cover net with an 8 mm mesh was set at the cod-end. All tows were carried out during the daytime at a mean ship speed of 3.0 knots for 30 min. The tow area (i.e., survey effort) of each station was calculated by recording the arrival and departure point on the bottom and horizontal open width of the net using the Net Recorder system (Furuno Electric Co., Hyogo, Japan or Marport, Reykjavik, Iceland).

The caught crabs were divided into males and females, and the number of individuals was counted. For males, the *cw* and right (if not available, left) cheliped height was measured to the nearest 0.01 mm using a digital caliper (CD67-A20PM, Mitsutoyo, Kanagawa, Japan) to identify their maturity (Watson 1970; Fujita et al. 1988). For females, the *cw* was also measured and recorded, and the abdominal pleon was observed to determine maturity; adult females are characterized by a broad abdominal pleon after terminal molt (Yoshida 1941). Instars of snow crabs off Tohoku were distinguished by their *cw* intervals (Table 1) (Ueda et al. 2007). In snow crab, the number of molts after instar VI is once a year; this can be used as an age trait (Kuwahara et al. 1995). The minimal instar of the snow crab obtained by this scientific bottom trawl was instar VIII, where their interval of *cw* is 24–42 mm (Table 1).

**Table 1.** Relationships among carapace width, instar, terminal molting, and sex that decide if the crab is fishable or not.

| Sex | Instar categories in population model | CW intervals | Terminally molted | Fishable |
| --- | --- | --- | --- | --- |
| Male | VIII | 24 - 42 | No | No |
|  | IX | 42 - 56 | No | No |
|  | X with not TM | 56 - 74 | No | No |
|  | X with TM | 56 - 74 | Yes | No |
|  | XI with not TM and not fishable | 74 - 80 | No | No |
|  | XI with TM and not fishable | 74 - 80 | Yes | No |
|  | XI with not TM and fishable | 80 - 86 | No | Yes |
|  | XI with TM and fishable | 80 - 86 | Yes | Yes |
|  | XII with not TM | 86 - 98 | No | Yes |
|  | XII with TM | 86 - 98 | Yes | Yes |
|  | XIII with not TM | 98 - | No | Yes |
|  | XIII with TM | 98 - 110 | Yes | Yes |
|  | XIV with TM | 110- | Yes | Yes |
| Female | VIII | 24 - 42 | No | No |
|  | IX | 42 - 56 | No | No |
|  | X with not TM | 56 - 74 | No | No |
|  | XI with TM and fishable | - | Yes | Yes |

We then calculated the density at the station and estimated the total number at instar (*na*) with their coefficients of variation (CVs) for the whole Tohoku region by multiplying by the area based on a swept area method (Shibata et al. 2019). CVs were corrected by Taylor’s power law (Supporting Information 1). The estimated catch efficiency (Hattori et al. 2014) of the trawl net and their variance–covariance matrix were used to estimate an unbiased abundance (see below eq. [34]).

### Catch data

Snow crab have only been caught off the coast of the Tohoku region by offshore bottom trawl fisheries (> 17 gross ton); their annual catches of snow crab were therefore used as a total catch. The catch statistics were distinguished for males (only for *cw* ≥ 80mm) and females (only for matured). Crabs were sampled and their *cw* measured to reflect a whole composition of *cw* in the total catch by the Fukushima Fisheries Experimental Station. Additionally, their right or left cheliped height (only for male) and maturity (only for female) were measured. Then, catch at instar (*ca*) was calculated based on the instar and maturity composition in the sample data. Because Fukushima Prefecture occupied 78% of fish catches (on average) of the total catch during 1997–2018 (Figure 2), the representativeness of the samples was guaranteed. Samples during 1997–1998 and 2003 were not available because the sample sizes were small, therefore the instar and maturity composition in 1999 and an average composition of 2002 and 2004 were used, respectively. Sample data during 2008–2010 were also not available because a hard disk containing those data was lost to the massive tsunami in March 2011. Therefore, a composition of 2007 data was used to obtain *ca* values for those years. Because there were few catches and measurements were not carried out from 2011 to 2017, compositions of this period were substituted by those of the scientific bottom trawl survey. In 2018, *ca* was available because measurements were carried out by scientists working for the FRA and Fukushima Prefectural Research Institute of Fisheries Resources.

### Statistical modeling

We developed a state-space stock assessment model coupled with an RW process, such as fishing mortality coefficient (*F*) used in the SAM (Nielsen and Berg 2014). We hereafter refer to it as “just another state-space stock assessment model” (JASAM). In JASAM, *M* and *p* can be stably estimated because the number at instar and maturity have been obtained based on the scientific bottom trawl survey. JASAM has two model structures, state and observation models, for the modeling of latent population dynamics and the catch (observation) process. Unlike in SAM, not only *F*, but also *M* and *p*, can have RW schemes in this model (see below).

### Modeling of natural mortality coefficient

#### Definition of M at group

There are six instar categories, where *a* (*a* = 8,…, 13) shows the instar. Although we are interested in an instar-specific natural mortality rate *M_a_*, the adjacent instars may take the same *M* because individuals of adjacent instars have similar body sizes, habitats, and are exposed to similar environments. In this study, instar group *g* (*g* = 0, …,5) was used to select *M* at group (*M_g_*) and instars were categorized for all groups (Supporting Information 2). The group consisted of successive instar groups. For example, if instar VIII and X are in the same group, instar IX is also in the same group. Since there are five “partitions” between one and six as integers and the same group cannot be formed across the partitions, 2^5^ = 32 combinations were made for *M_g_*.

#### Time-varying M

As we explained in the introduction, it was suspected that *M* and/or *p* have maintained a high value in recent years. Considering the possibility that *M* varies with time, variations in *M* were assumed to be one of the following three patterns: a constant, an RW of first-order difference, or that of second-order difference. In the constant style, *M_g_*_,*t*+1_ = *M_g_*_,*t*_ = *M_g_* where *t* shows year (*t* = 1997,…,2018). In the first-order RW style,

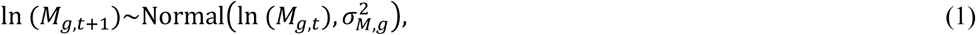

where *σ_M_* is the standard deviation of the normal distribution used for RW. In the second-order RW style, this is shown as

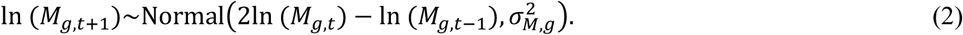

These three patterns of *M_g,t_* were selected by model selection (see below).

#### M after the terminal molting

We categorize the number of years elapsed after the terminal molting, *j* (*j* = 0, 1, 2), into immature (*j* = 0), terminally molted within one year (*j* = 1), and terminally molted after more than one year (*j* = 2) (Shibata et al. 2019). The crabs mature functionally at the same time that they undergo the terminal molt and cease to grow. Then, since the shells of snow crab have started to harden gradually, *M* can be lower than that of an immature crab (Yamasaki 1988; Yamasaki et al. 1992). In the stock assessment of snow crab, it has been assumed that *M* decreases for individuals one year after terminal molting (*j* = 2). Therefore, *M* of an individual one year after the terminal molt was multiplied by a multiplier *φ* (0 < *φ* <1) to express the change in *M* after the terminal molt:

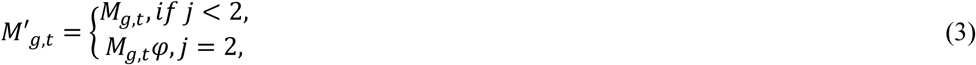

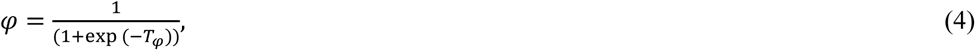

where *M_g_*_,*t*_′ is the natural mortality rate corresponding to the number of years elapsed after the terminal molting and *T_φ_* is a parameter that should be estimated.

### Modeling of fishing mortality coefficient

Because the spatial distribution of snow crab off Tohoku is basically divided between mature and immature individuals by sex rather than instar (Kitagawa 2000), *F* at instar has not been estimated in the stock assessment in Japan (Shibata et al. 2019). We also use fishing mortality *F*, specified by maturity status and sex. The time-varying *F* is shown below:

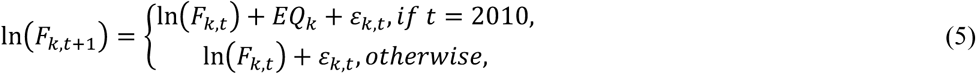

where *k* = 1 (immature male), *k* = 2 (mature male), and *k* = 3 (mature female). When *t* is 2010, the rapid decrease in fishing pressure due to the earthquake cannot be expressed by RW; therefore, it is estimated as a fixed effect *EQ*. Here,

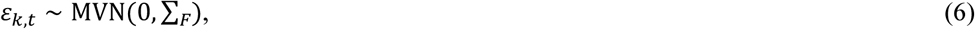

where *ε* follows a multivariate normal (MVN) distribution, and its variance–covariance matrix Σ*_F_* is shown as

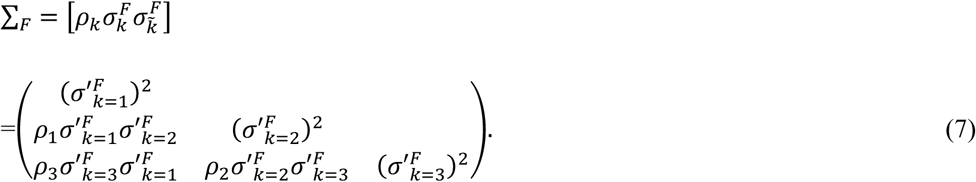

Here, upper triangular components were omitted and *ρ* and *σ′* were changed after 2011 as below:

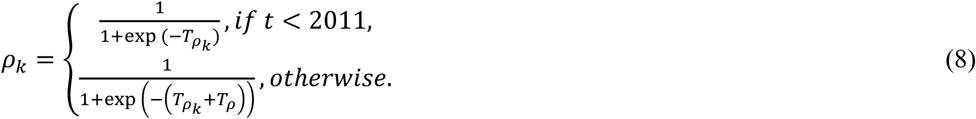

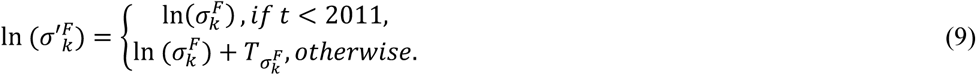

Here, *T_ρ_* is tested for whether it is zero in a model selection (see below), although *T_σ_* is not, because the total catch has apparently decreased and therefore *T_σ_* must be changed after 2011. Model selection was also carried out for *ρ_k_* in five cases: one case that all *ρ_k_* are one (i.e., *T_ρk_* is not estimated), three cases that two of the three *ρ_k_* take the same value, and one case that all *ρ_k_* are different.

### Modeling of terminal molt probability

The terminal molt probability (*p*) was modeled as a function of the instar; *p* may vary with time. We modeled *p* using an RW process, as below:

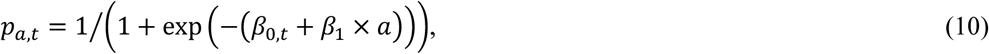

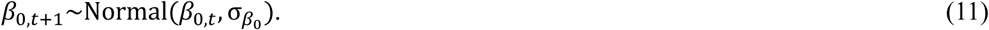

We carried out model selection to determine whether *β*_0,*t*_ = *β*_0_ (i.e., not time-varying but constant) or *β*_0,*t*_.

### State model of male

The structure of snow crab population dynamics is quite complex because they have six-plus groups after instar X and the male fishable size (*cw* ≥ 80 mm) divided instar XI (*cw* interval is 74–86 mm) into two categories (Table 1). We hereafter describe the model formulae of each transition step by step. Here, the initial number (i.e., in *t* = 1997) of snow crab at instar and sex were parameters to be estimated.

#### From instar VIII to IX (a = 8)

The number at instar from instar VIII to IX can be shown as below:

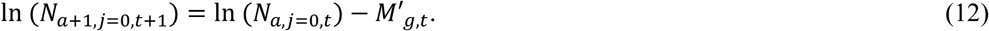

#### From instar IX to X (a = 9, immature)

From instar IX to X, some individuals mature (undergo terminal molting). The population dynamics model can be expressed as

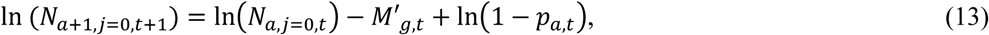

where 1 – *p_a_*_,*t*_ is the probability that an individual is not terminally molted.

#### From instar IX to X (a = 9, mature)

Terminal molting at instar X results in maturing with a *cw* of less than 80 mm, and the individual ends its life without recruiting to a stock. The dynamics from immature to mature are shown as

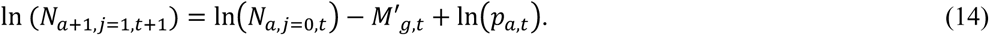

A plus group of instar X is shown as below:

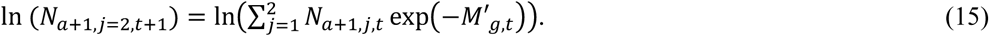

#### From instar X to XI (a = 10, immature)

Since male snow crabs with a *cw* larger than 80 mm are fishable, the number of crabs at instar XI (74–86 mm) multiplied by *r* (0 < *r* < 1) are fishable. In other words, the number of crabs at instar XI were separated into ranges (74–80 mm as not fishable and 80–86 mm as fishable). Although we had assumed that *r* was 0.5 in a previous study (Shibata et al. 2019), we estimated *r* in this report. Individuals of instar XI with a *cw* of 74–80 mm without terminal molting can be modeled as

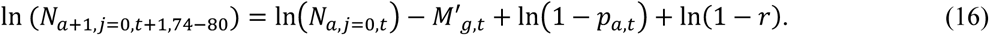

The individuals of instar XI with a *cw* of 80–86 mm without terminal molting is then modeled as

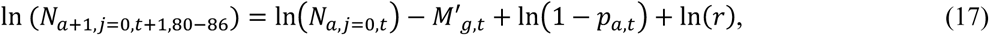

where *r* = 1/(1 + exp(–*T_r_*)) and *N_a_*_+1,*j*=0,*t*+1_ = *N_a_*_+1,*j*=0,*t*+1,74−80_ + *N_a_*_+1,*j*=0,*t*+1,80−86_.

#### From instar X to XI (a = 10, mature)

Individuals of instar XI with a *cw* of 74–80 mm with terminal molting are modeled as

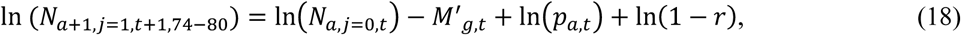

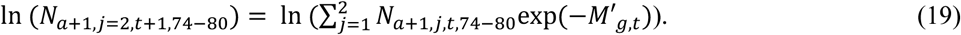

Individuals of instar XI with a *cw* of 80–86 mm with terminal molting are modeled as

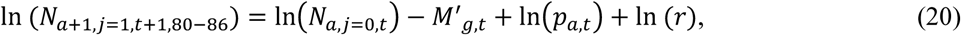

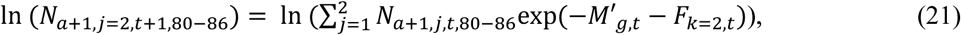

where individuals that had experienced terminal molting were caught with a fishing mortality coefficient of *F*.

#### From instar XI to XII (a = 11, immature)

Of the 74–80 mm and 80–86 mm individuals at instar XI, only the latter is subject to catch and is expressed as follows:

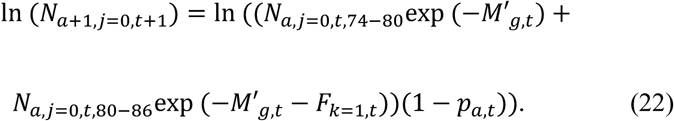

#### From instar XI to XII (a = 11, mature)

Because the 80–86 mm individuals are terminally molted and included into a plus group of instars as

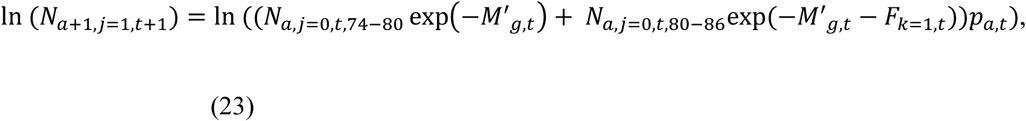

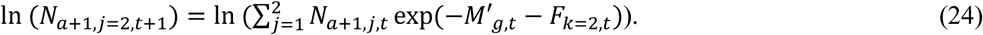

#### From instar XII to XIII (a = 12, immature)

The number at instar from instar XII to XIII not terminally molted can be shown as below:

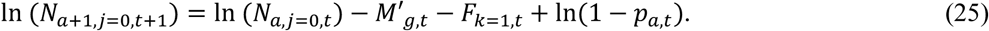

#### From instar XII to XIII (a = 12, mature)

The number at instar from instar XII to XIII terminally molted can be shown as below:

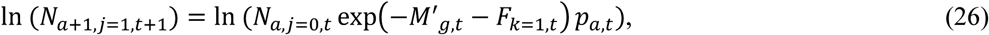

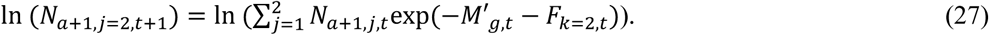

#### From instar XIII to XIV (a = 13)

Because all individuals at this instar stage mature at probability one, the equation is shown as below:

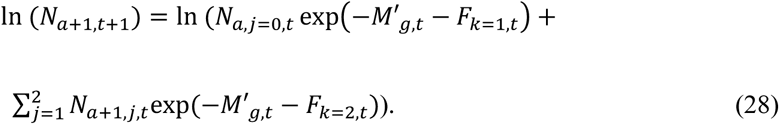

### State model of female

#### From instar VIII to IX and IX to X (a = 8 and 9, immature)

Because females do not mature at instar X, the population transition is described as

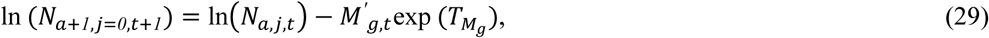

where *T_Mg_* is the female-specific term, although *T_Mg_* is tested for whether the term is zero or not.

#### From instar X to XI (a = 10, immature)

In females, because only instar XI is fishable, the transition is shown as below:

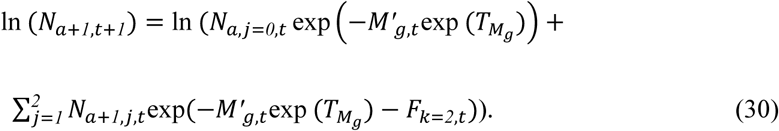

#### Estimation of the number of individuals at instar VIII

Because our survey can observe snow crabs older than instar VIII, we treated the number of individuals at instar VIII as recruits. Here, we assumed an RW process as below:

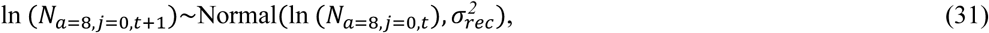

where the numbers of males and females were assumed to be the same (i.e., the sex ratio of recruits was assumed as 0.5).

### Observation model

#### Scientific bottom trawl survey

The estimated number of individuals in the trawl survey *n* is obtained by multiplying the catch efficiency *q* by the true number of individuals *N*. Elapsed years after the terminal molting are not known by the trawl survey; only the identification before or after the terminal molting (*u* = 0 where *j* = 0, *u* = 1 where *j* = 1 and 2) can be determined. An observation model for the trawl survey is shown as below:

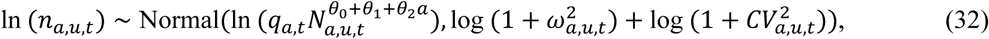

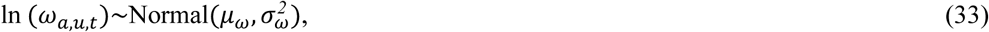

where *θ*_0_–*θ*_2_ are parameters of hyperstability (or hyperdepletion) that show a nonlinear relationship between abundance and its index (Hilborn and Walters 1992; Chen et al. 2008), and *θ*_1_ is only estimated for females. To make the model flexible, we treated *ω*^2^ as a random effect term. We calculated the likelihood as both *n_a,u,t,_*_74–80_ and *n_a,u,t,_*_80–86_ for males, although the suffix was omitted in eq. (32). The CV of the number of individuals estimated by the swept area method is used in eq. (32). The catch efficiency *q* is shown as below:

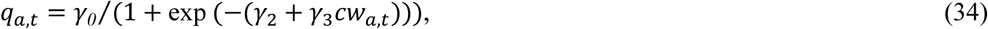

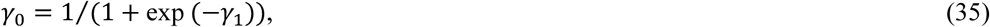

where *cw_a_*_,*t*_ was the average *cw* of each instar obtained from the annual trawl survey. The catch efficiency *q_a,t_* was treated as a random effect term and the average *γ*_1_–*γ*_3_ and their variance–covariance matrix was plugged in from the previous study (Hattori et al., 2014).

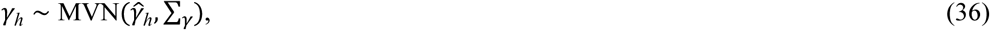

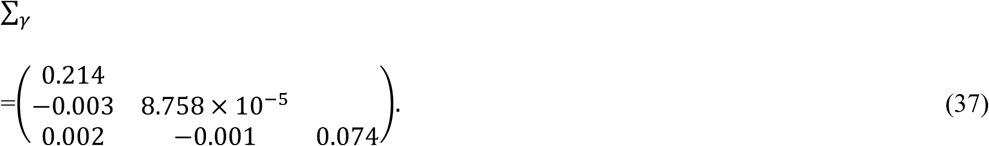

Here, upper triangular components were omitted and 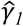 = 0.683, 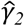 = −4.276, and 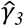 = 0.0792.

#### Catch at instar

*c_a_* is the observed number of catch at instar and *C_a_* is the estimated number of catch at instar; these were shown as below:

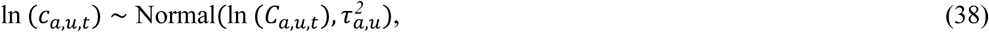

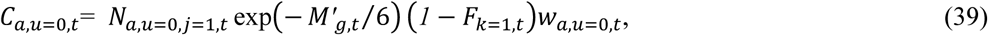

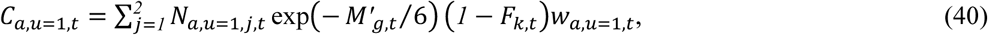

where the catch of male snow crab was applied using both eq. (39) and (40) (*k* = 2), although that of females was applied using eq. (40) (*k* = 3) alone, because only mature females were caught.

### Model selection

#### State and observation models

There are eight factors to be selected in the model: 1) The variables/types of difference in *M_g_* to be selected were 32 combinations for *M_g_* (Supporting Information 2), 2) three types of difference for *M_g,t_*, 3, 4) either *φ* and *T_Mg_* = 1 or not, 5) either *T_ρ_* = 0 or not, 6) five combinations for *T_ρk_*, 7) either *β*_0_ was time-varying or not, and 8) either the parameters of hyperstability were included in model or not (Table 2). Consequently, the number of tested models is 15,360 (=32 × 3 × 2 × 2 × 2 × 5 × 2 × 2). The number of parameters for *M_g,t_* were all about whether a constant (*M_g_*), a first-order difference (*σ_M_*^2^*_g_*), or a second-order difference (*σ_M_*^2^*_g_*) model was selected, because the number of parameters equaled the number of groups. The first-order difference may be selected more easily than the other two types because it is considered to be the most flexible for fitting. We therefore performed model selection for each of three types for *M_g,t_* and selected the best one by both AIC (Akaike 1974) and Bayes’ information criterion (BIC) (Schwarz 1978). Six models will ultimately be chosen as candidates for the best model through this procedure.

**Table 2.**
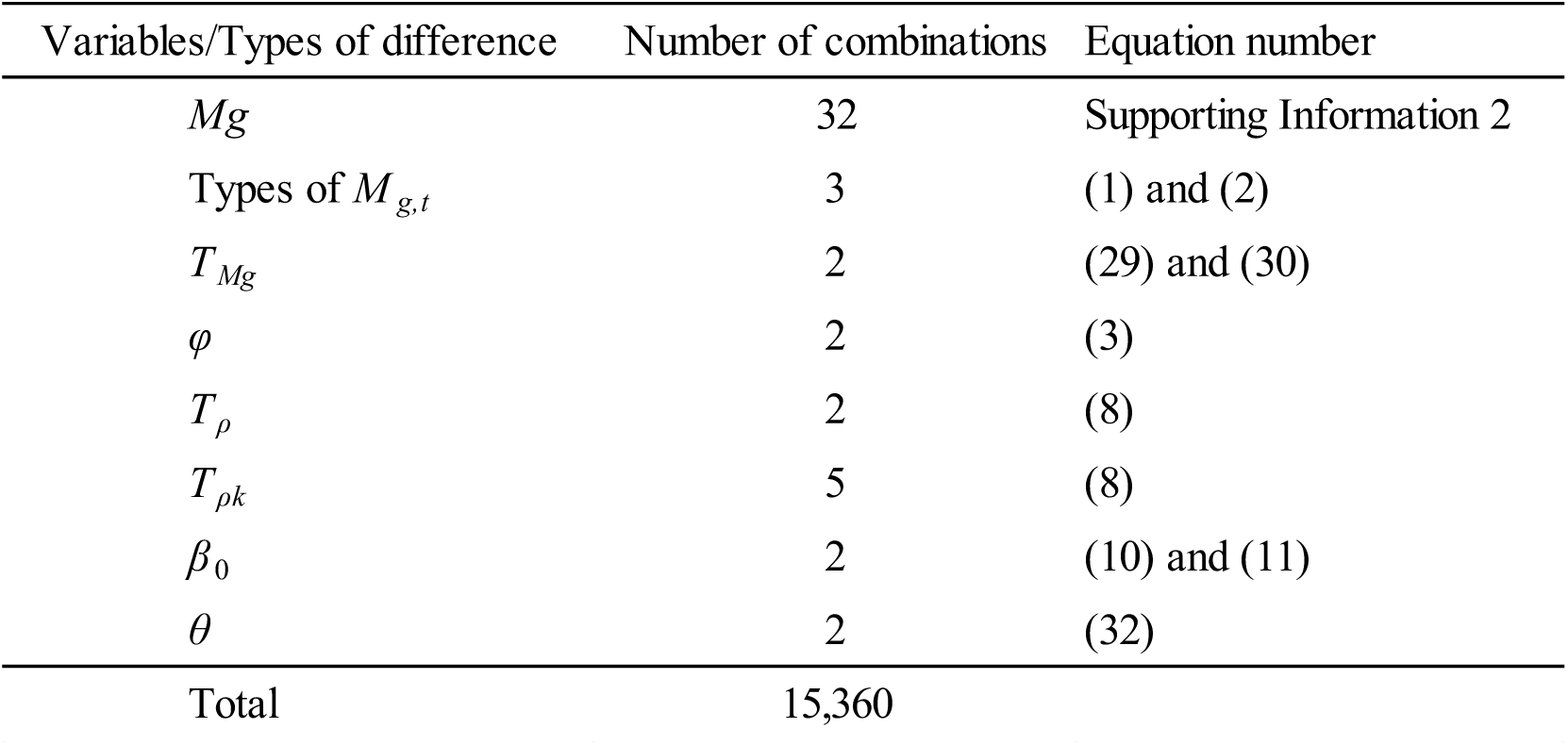
Combination of all variables/types of difference for model selection.

| Variables/Types of difference | Number of combinations | Equation number |
| --- | --- | --- |
| $Mg$ | 32 | Supporting Information 2 |
| Types of $M_{g,t}$ | 3 | (1) and (2) |
| $T_{Mg}$ | 2 | (29) and (30) |
| $\varphi$ | 2 | (3) |
| $T_{\rho}$ | 2 | (8) |
| $T_{\rho k}$ | 5 | (8) |
| $\beta_0$ | 2 | (10) and (11) |
| $\theta$ | 2 | (32) |
| Total | 15,360 |  |

The estimated abundance *A_T−i_*(*A* = ∑*_a_N_a_w_a_*, *T* = 2018, *i* = 1,…, 5) of both males and females was calculated using all the data from 1997 to 2018. The estimated abundance using the data period from 1997 to *T* – *i* (*i* = 1,…, 5) was denoted as 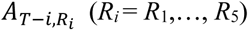, where *R_i_* is a suffix indicating how many years of data are excluded. As an index representing retrospective bias, Mohn’s rho (*ρ_past_*) (Mohn 1999) was calculated by the following equation:

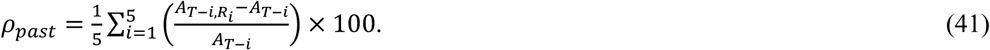

In addition, a retrospective forecasting (Brooks and Legault 2015) approach was used to estimate an error in future projections for two years ahead because the ABC of snow crab had been calculated based on data from two years ago (Shibata et al. 2019). First, the abundance was estimated using the data from 1997 to *T* – *j*, excluding the data for *j* years (*j* = 3,…, 7). Then 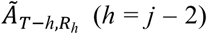 was projected for the abundance two years ahead. For example, when *j* = 3 then *h* = 1, the abundance and the parameter estimation were performed using the data up to 2015, and the abundance in 2017 was predicted. This procedure was repeated to calculate *ρ_future_* for ABC using the following formula:

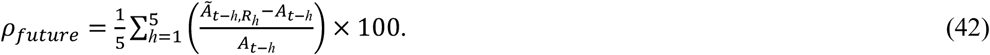

All parameter values were given their average over the past three years to calculate *ρ_future_*. Biases in the estimation and prediction of the abundance for the six best models were evaluated using *ρ_past_* and *ρ_future_* and the best model was selected. As a sensitivity test, the CVs of the observed number of instars were multiplied by 1.5 using the best model. We then calculated *ρ_past_*, *ρ_future_*, and time-varying *M_g,t_*.

#### Stock–recruitment relationship

We fitted three types of stock–recruitment (SR) relationship between the estimated spawning stock biomass (SSB) and recruitment (instar VIII) that were obtained from the best model from the above model selection phase for future predictions. We used three types of SR relationships as hockey stick (HS) (Clark et al. 1985), Beverton–Holt (Beverton and Holt 1957) (BH) and Ricker (RI) (Ricker 1954) models for eq. (43), (44), and (45), respectively, as below:

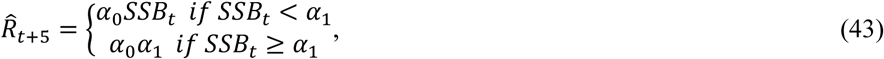

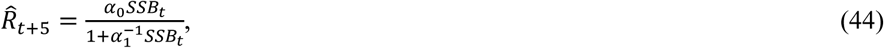

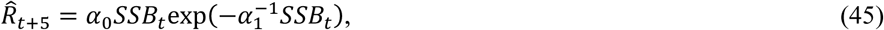

where *t* is year (*t* = 1997,…,2013). In snow crab, although there is no information on the length of each instar duration in the Tohoku region, it has been assumed that five years are needed to reach instar VIII in the Sea of Japan (Ueda et al. 2007), and we assumed this to be true in these equations. SSB is the spawning stock biomass after a fishing season and is calculated as below:

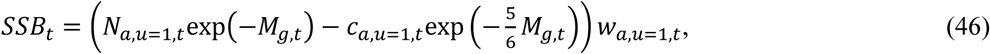

where *N*, *c*, and *w* are the estimated number, observed catch number, and mean weight of mature female, respectively. *α*_0_ and *α*_1_ are parameters to be estimated by maximizing the log likelihood (*LL*) function for each model, as below:

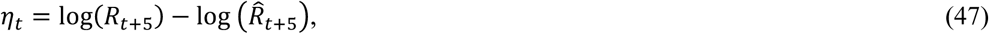

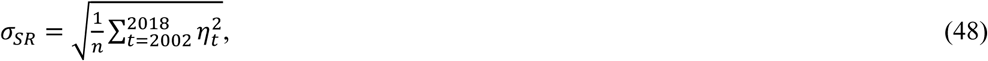

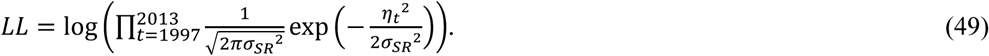

Snow crab could have a warm and cold regime for their recruitment in the eastern Bering Sea (Szuwalski and Punt 2013). Although it has not been reported surrounding Japan, we considered the case that an autocorrelation existed in residuals (*η*) to express the regime of recruitment, as shown below:

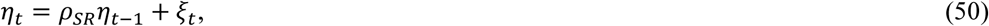

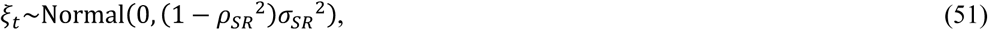

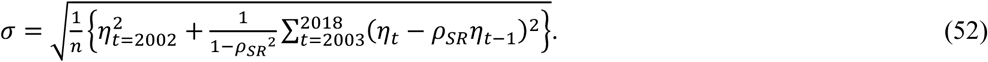

Here, we did not estimate *α*_0_, *α*_1_, and *ρ_SR_* simultaneously because it had been reported that estimates would be unstable and bias could arise (Johnson et al. 2016). In summary, we carried out a model selection using AICc (Hurvich and Tsai 1989) from the three SR relationships (i.e., *α*_0_ and *α*_1_ were fixed). We then estimated *ρ_SR_* and tested whether the autocorrelation in the residuals estimated for *ρ_SR_* was zero or not.

### Estimation of maximum sustainable yield

Because *M* and *p* were time-varying, we defined their values used to estimate the MSY. We prepared three scenarios in *M* and *p* as 1) the mean values during 2016–2018, 2) mean values during 1997–1999, and 3) mean values among all years. In the future prediction, the best model of the SR relationship was used. To estimate the MSY, we used the below equations for *F*:

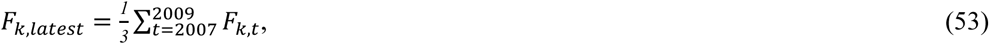

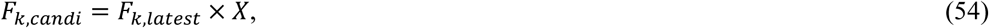

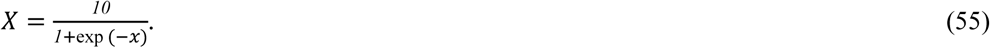

We changed *x* from –10 to 10 by 0.01 and *F_k_*_,*candi*_ was used for the future prediction. The life expectancy of snow crab in Newfoundland was reported to be 13 and 19 years old for females and males, respectively (Comeau et al. 1998). Although it has not been studied in detail in Japan, snow crab life expectancy was often assumed to be more than 10 years old (e.g., Shibata et al., 2019). We assumed the life span of snow crab to be 15 years and simulated the future prediction as 20 times the life span to obtain initial values of the population. We carried out the prediction for 400 years and the mean catches between 301 and 400 years were recorded where the first 300 years were not used to delete the effects of the initial values. We repeated this procedure 1,000 times and calculated a median catch each *x* (i.e., a mean catch between 301 and 400 years was obtained 1,000 times each *x* and 2,001 medians from the mean catches were obtained). Then, we selected the *F_k_*_,*candi*_ that maximized the medians of catch as *F_MSY_*, and MSY and SSB_MSY_ were also obtained. The calculation was carried out using freely available statistical analysis software R (R Core Team 2019) and the Template Model Builder (TMB) (Kristensen et al. 2015).

## Results

### The best models of state and observation models

Models that had minimal AIC and BIC values are shown in Table 3. The result also showed the combination of *g*, the variables included in a model, and values of retrospective analysis for each model. The models with constant *M_g_* through time were the same as each other regardless of the two information criteria. The model showed that instars VIII and IX, and instars X, XI, and XII were the same groups. The models with a first-order difference of *M_g_* were the same whether the criterion was AIC or BIC where instars VIII and IX, instars X and XI, and instars from XII to XIII were grouped, respectively. The parameter *T_Mg_* was selected in the model. The model had both the minimum AIC (1,064.5) and BIC (1,254.2) among the three formulations of *M_g,t_*. In the case of the model with a second-order difference of *M_g_*, both AIC and BIC had the smallest values when all of the age groups were combined. In contrast, although the AIC minimal model contained three parameters (*θ*_0_–*θ*_2_) that considered hyperstability, the BIC minimal model did not. The parameter of terminal molting probability *β* was selected as time-varying, and *T_ρk_*_=1_ was not different from *T_ρk_*_=2_ in all cases.

**Table 3.** Results of the model selection and retrospective analysis. The column *M* showed three types of *M_g,t_* (constant, first-order, or second-order difference). *M_g_* was same if the numbers expressed in columns from VIII to XIII were the same. If the cell showed “In”, that indicated that the variable was selected in the model by information criteria, and “-” showed that it was not selected. *β* was selected as *p* was time-varying (indicated as “*t*”) in all cases.

| *M* | *Criterion* | *VIII* | *IX* | *X* | *XI* | *XII* | *XIII* | $T_{\varphi}$ | $T_{Mg}$ | $T_{\rho}$ | $\theta$ | $\beta$ | AIC/BIC | $\rho_{past}$ | $\rho_{future}$ |
| --- | --- | --- | --- | --- | --- | --- | --- | --- | --- | --- | --- | --- | --- | --- | --- |
| Const. | AIC | 0 | 0 | 1 | 1 | 1 | 2 | - | - | - | In | *t* | 1147.3 | -5.6 | -8.5 |
|  | BIC |  |  |  |  |  |  |  |  |  |  |  | 1345.2 |  |  |
| First | AIC | 0 | 0 | 1 | 1 | 2 | 2 | - | In | - | In | *t* | 1064.5 | 0.2 | 66.3 |
|  | BIC |  |  |  |  |  |  |  |  |  |  |  | 1254.2 |  |  |
| Second | AIC | 0 | 0 | 0 | 0 | 0 | 0 | - | - | - | In | *t* | 1118.2 | -12.5 | -43.3 |
|  | BIC | 0 | 0 | 0 | 0 | 0 | 0 | - | - | - | - | *t* | 1311.4 | 1.2 | 3.1 |

The values of *ρ_past_* did not greatly change among models and were relatively small (Table 3). In contrast, *ρ_future_* showed poor performance except for in the BIC best model with a second-order difference of *M_g_*_,*t*_. Although the model with a first-order difference of *M_g,t_* had the smallest AIC and BIC among the three formulations of *M_g,t_*, we decided that the minimal BIC model with a second-order difference of *M_g,t_* was the best model from the synthetic evaluation of model performance, including retrospective bias and retrospective forecasting. All of the estimated parameter values of the best model are shown in Table 4, and the results indicated that the estimated values were well fitted to the observed values and the residuals showed normally distributed (Figure 3 and Supporting Information 3)

**Fig. 3.**
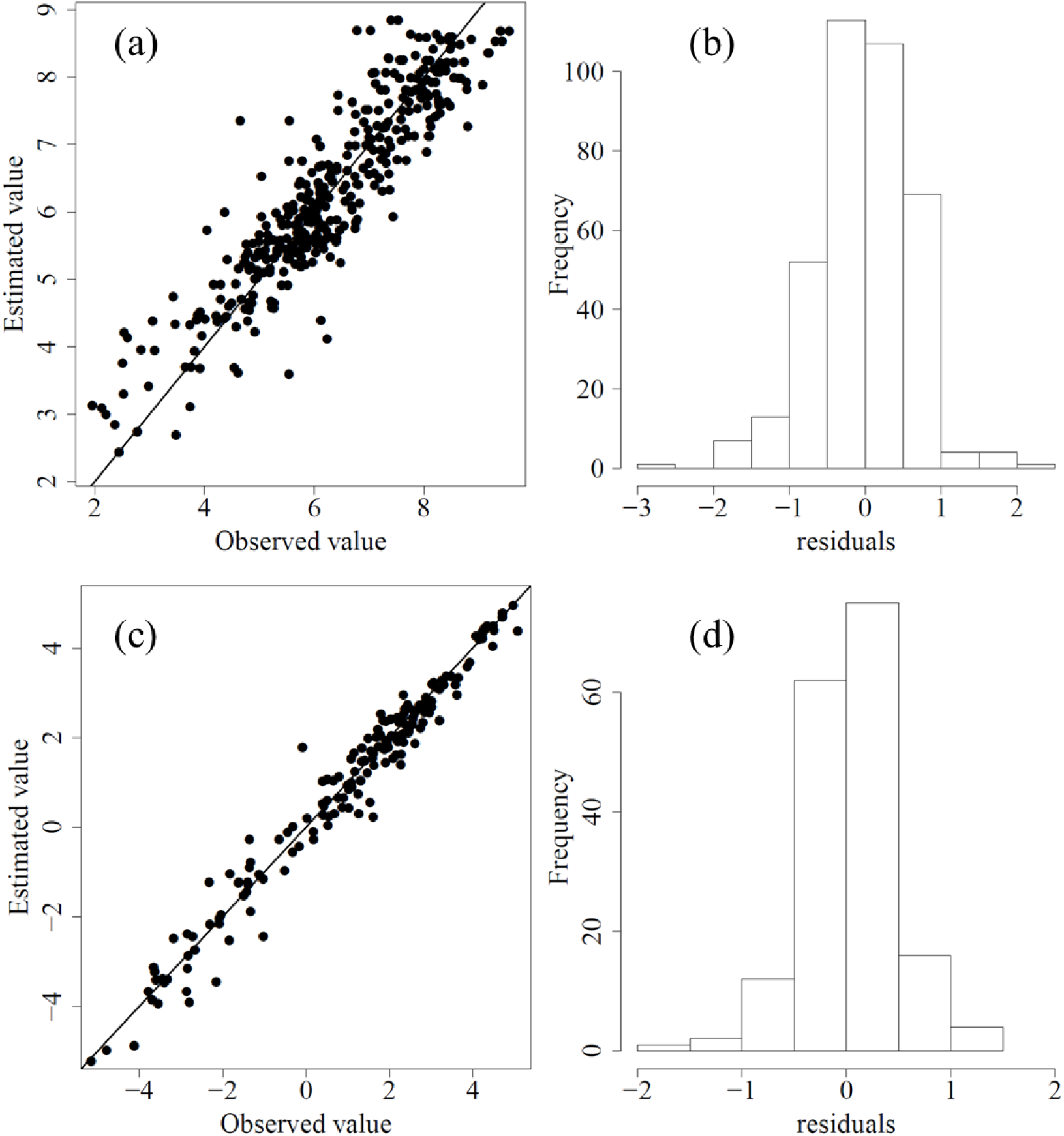
Relationship between observed and estimated values and histograms of residuals for the number of snow crabs (a, b) and catch (c, d) of the best model.

**Table 4.** Estimated parameter values and their standard errors by the best second-order difference model.

| Parameters | Estimates | Std.error | Either male or female |
| --- | --- | --- | --- |
| $\ln((\sigma_{k=1}^F)^2)$ | -0.170 | 0.239 | male |
| $\ln((\sigma_{k=2}^F)^2)$ | -1.349 | 0.367 | male |
| $\ln((\sigma_{k=3}^F)^2)$ | -1.491 | 0.725 | female |
| $T_{\sigma}^F_{k=1}$ | 0.660 | 0.368 | male |
| $T_{\sigma}^F_{k=2}$ | 1.829 | 0.474 | male |
| $T_{\sigma}^F_{k=3}$ | 2.104 | 0.750 | female |
| $EQ_{k=1}$ | -5.020 | 0.948 | male |
| $EQ_{k=2}$ | -5.996 | 0.396 | male |
| $EQ_{k=3}$ | -5.540 | 0.403 | female |
| $\ln(\sigma_{M,g}^2)$ | -2.998 | 0.771 | both |
| $\ln(\tau_{a=11,u=0}^2)$ | -0.994 | 0.467 | male |
| $\ln(\tau_{a=11,u=1}^2)$ | -0.934 | 0.220 | male |
| $\ln(\tau_{a=12,u=0}^2)$ | -0.787 | 0.329 | male |
| $\ln(\tau_{a=12,u=1}^2)$ | -1.076 | 0.209 | male |
| $\ln(\tau_{a=13,u=0}^2)$ | -0.221 | 0.210 | male |
| $\ln(\tau_{a=13,u=1}^2)$ | -1.100 | 0.231 | male |
| $\ln(\tau_{a=14,u=1}^2)$ | -0.332 | 0.166 | male |
| $\ln(\tau_{a=11,u=1}^2)$ | -1.299 | 0.372 | female |
| $\ln(N_{a=9,u=0,t=1997})$ | 6.969 | 0.243 | male |
| $\ln(N_{a=10,u=0,t=1997})$ | 6.335 | 0.243 | male |
| $\ln(N_{a=10,u=1,t=1997})$ | 4.916 | 0.561 | male |
| $\ln(N_{a=11,u=0,t=1997,74-80})$ | 5.352 | 0.352 | male |
| $\ln(N_{a=11,u=1,t=1997,74-80})$ | 4.450 | 0.466 | male |
| $\ln(N_{a=11,u=0,t=1997,80-86})$ | 5.023 | 0.350 | male |
| $\ln(N_{a=11,u=1,t=1997,80-86})$ | 5.410 | 0.278 | male |
| $\ln(N_{a=12,u=0,t=1997})$ | 4.714 | 0.342 | male |
| $\ln(N_{a=12,u=1,t=1997})$ | 5.318 | 0.274 | male |
| $\ln(N_{a=13,u=0,t=1997})$ | 3.308 | 0.586 | male |
| $\ln(N_{a=13,u=1,t=1997})$ | 5.525 | 0.245 | male |
| $\ln(N_{a=14,u=1,t=1997})$ | 4.937 | 0.300 | male |
| $\ln(N_{a=9,u=0,t=1997})$ | 6.794 | 0.461 | female |
| $\ln(N_{a=10,u=0,t=1997})$ | 7.613 | 0.492 | female |
| $\ln(N_{a=11,u=0,t=1997})$ | 8.295 | 0.345 | female |
| $\ln(\beta_1)$ | -0.060 | 0.060 | both |
| $\ln(\sigma_{\beta_0})$ | -1.596 | 0.677 | both |
| $T_{\rho,k=1}$ | 3.029 | 0.678 | both |
| $T_{\rho,k=3}$ | -1.179 | 0.312 | both |
| $T_r$ | -0.149 | 0.104 | both |
| $\ln(\sigma_{rec}^2)$ | -1.085 | 0.203 | both |
| $\mu_{\omega}$ | -0.803 | 0.089 | both |
| $\ln(\sigma_{\omega}^2)$ | -1.138 | 0.299 | both |

Estimated time-varying *M_g_*_,*t*_ and terminal molting probabilities from the best model were shown in Figures 4 and 5, respectively. Although the time-varying *M_g_*_,*t*_ was not so high in 1997 (*M_g_*_,*t*_ = 0.20), values increased from around 2005 to 2012. The *M_g_*_,*t*_ kept a high value of more than 0.59. This result indicated that the abundance of snow crab could not increase if the total catch was kept at quite low values after the earthquake because the natural mortality also kept a high value. Terminal molting probability also kept increasing from around 2005 in all instars that had terminal molts (Figure 5). Although the values were 0.09, 0.19, 0.38, and 0.61 in 1997 for each instar (IX, X XI and XII), these values increased to 0.21, 0.41, 0.64, and 0.82 in 2017. This indicated that the terminal molting probabilities increased 2.46-, 2.10-, 1.67-, and 1.34-fold, respectively from 1997 to 2018. This also showed that the decreasing abundance was caused by both the high natural mortality and terminal molt probability. The estimated abundance and SSB were shown in Figures 6 and 7, respectively. Both results showed that estimated values kept decreasing after 2008 prior to the earthquake when estimated fishing mortalities were kept quite low (*F* = 0.04 for immature male in 2016 was the maximum) after 2011 (Figure 8).

**Fig. 4.**
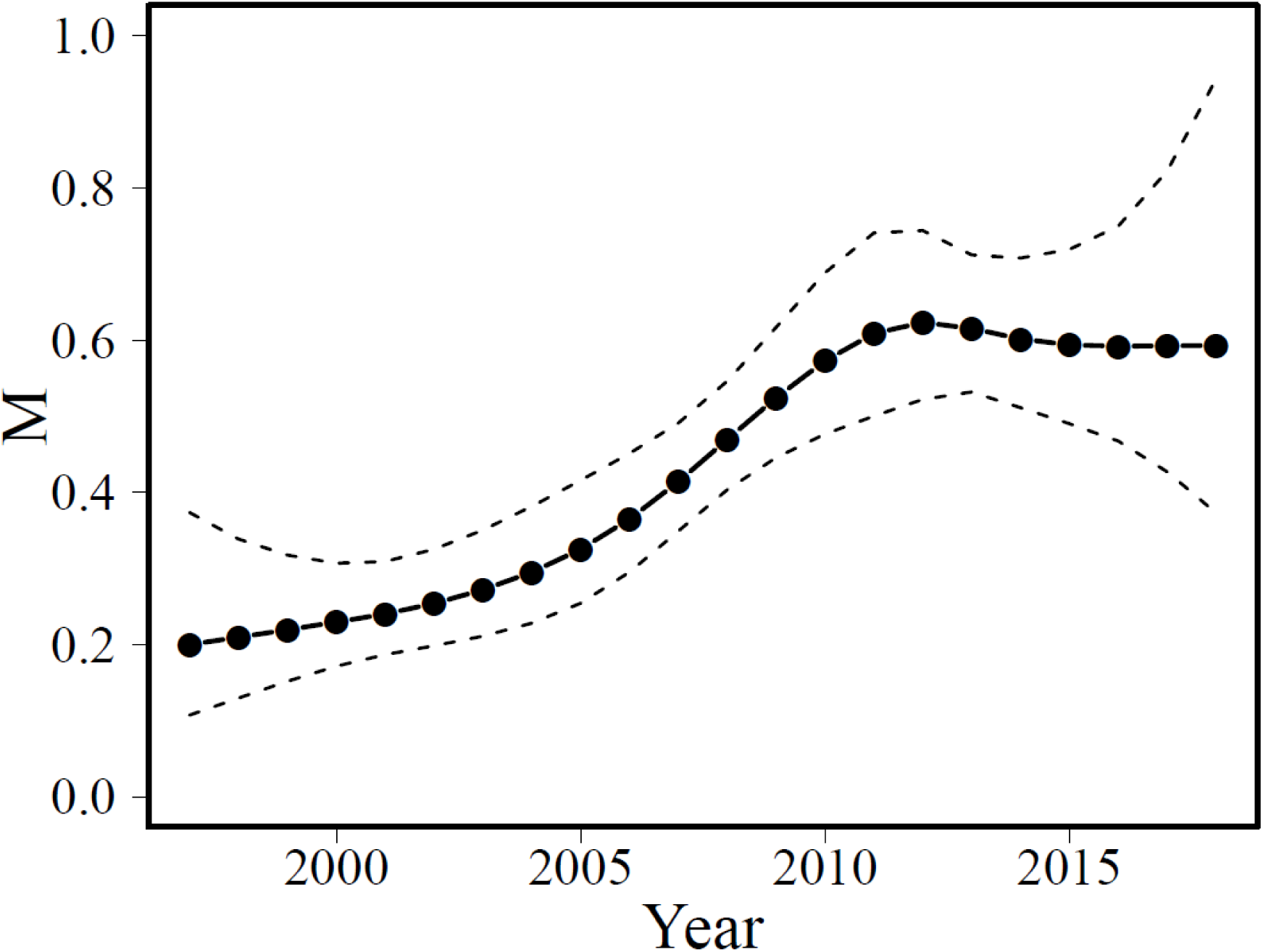
Estimated *M_g,t_* of the best model with a 95% confidential interval (break lines).

**Fig. 5.**
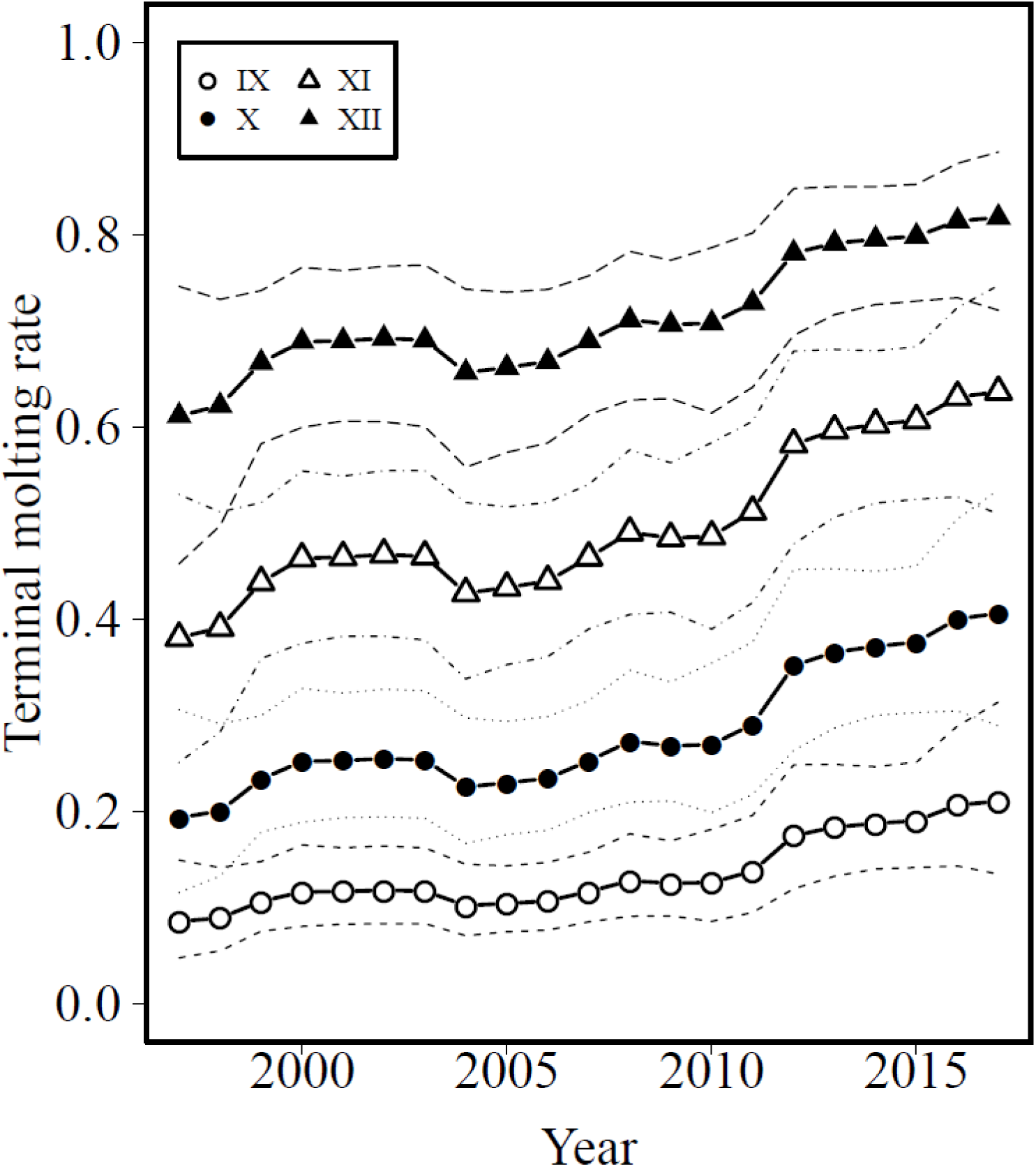
Estimated *p_a,t_* of the best model with their 95% confidential intervals (break lines). The symbols indicate those of instars IX (white circles), X (black circles), XI (white triangles), and XII (black triangles).

**Fig. 6.**
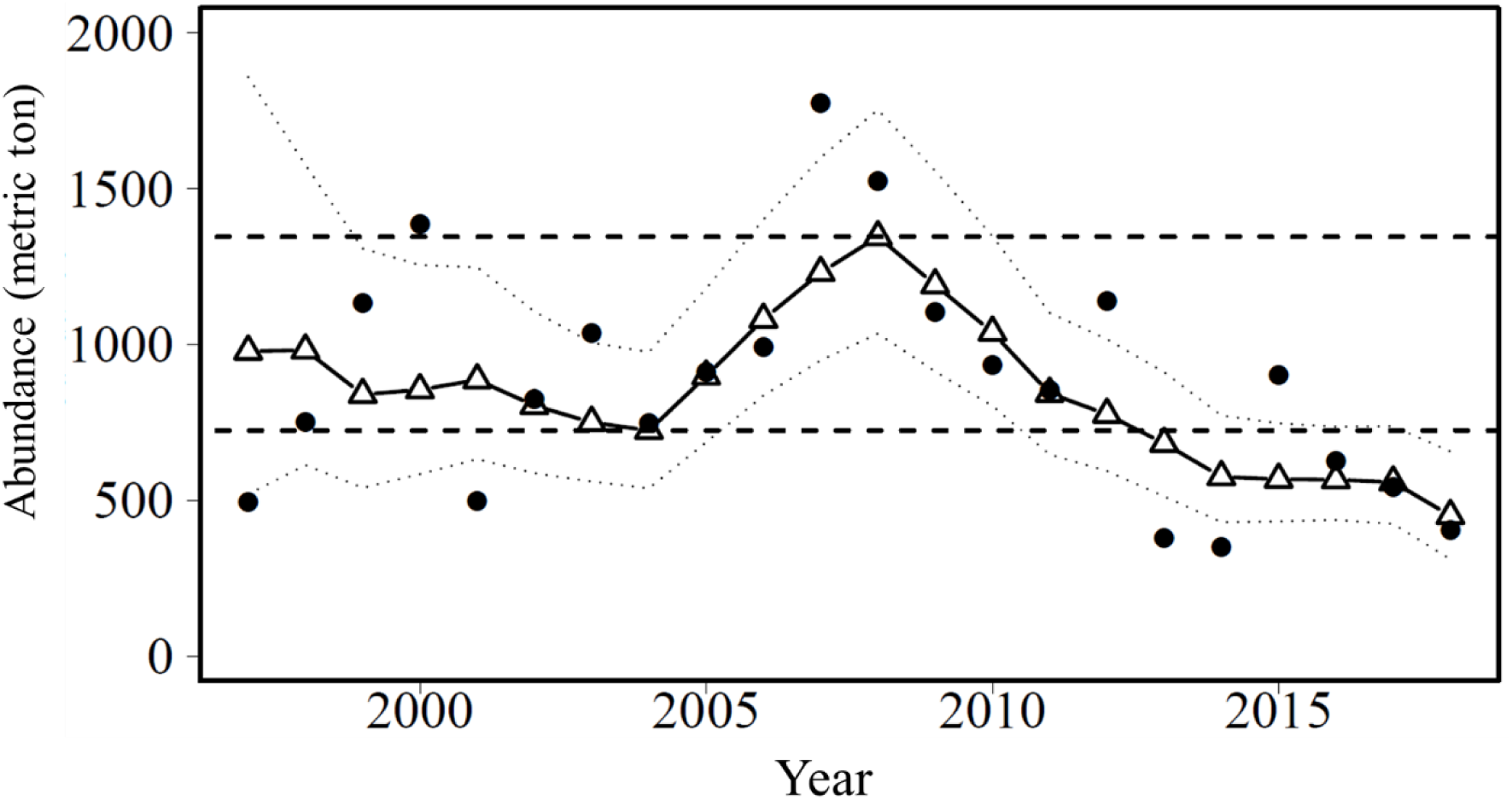
Estimated abundance of the best model with 95% confidential interval (break lines). The two horizontal dashed lines show the minimum and maximum estimates before 2011.

**Fig. 7.**
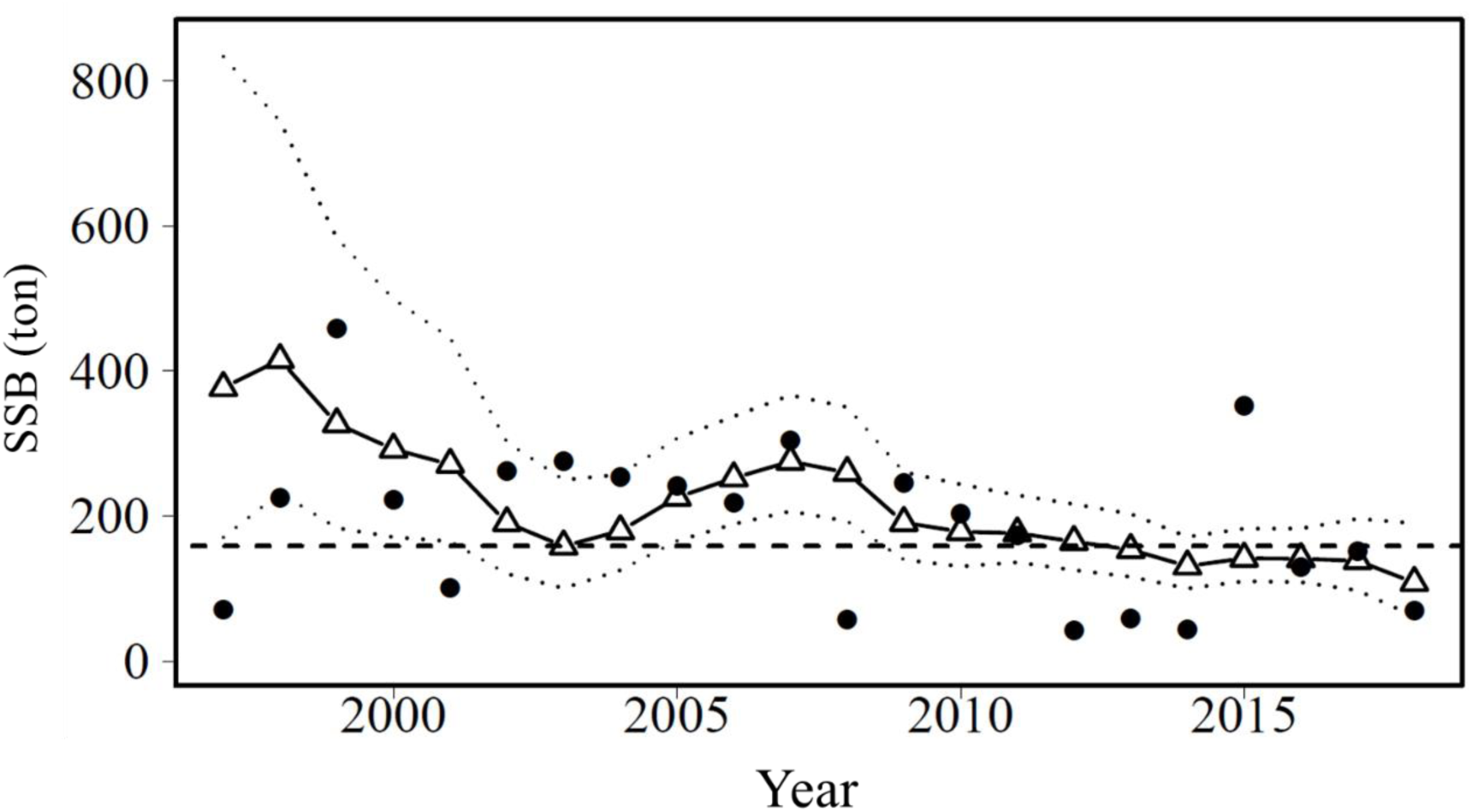
Estimated SSB after a fishing season of the best model with a 95% confidential interval (break lines). The horizontal dashed line shows the minimum estimates before 2011.

**Fig. 8.**
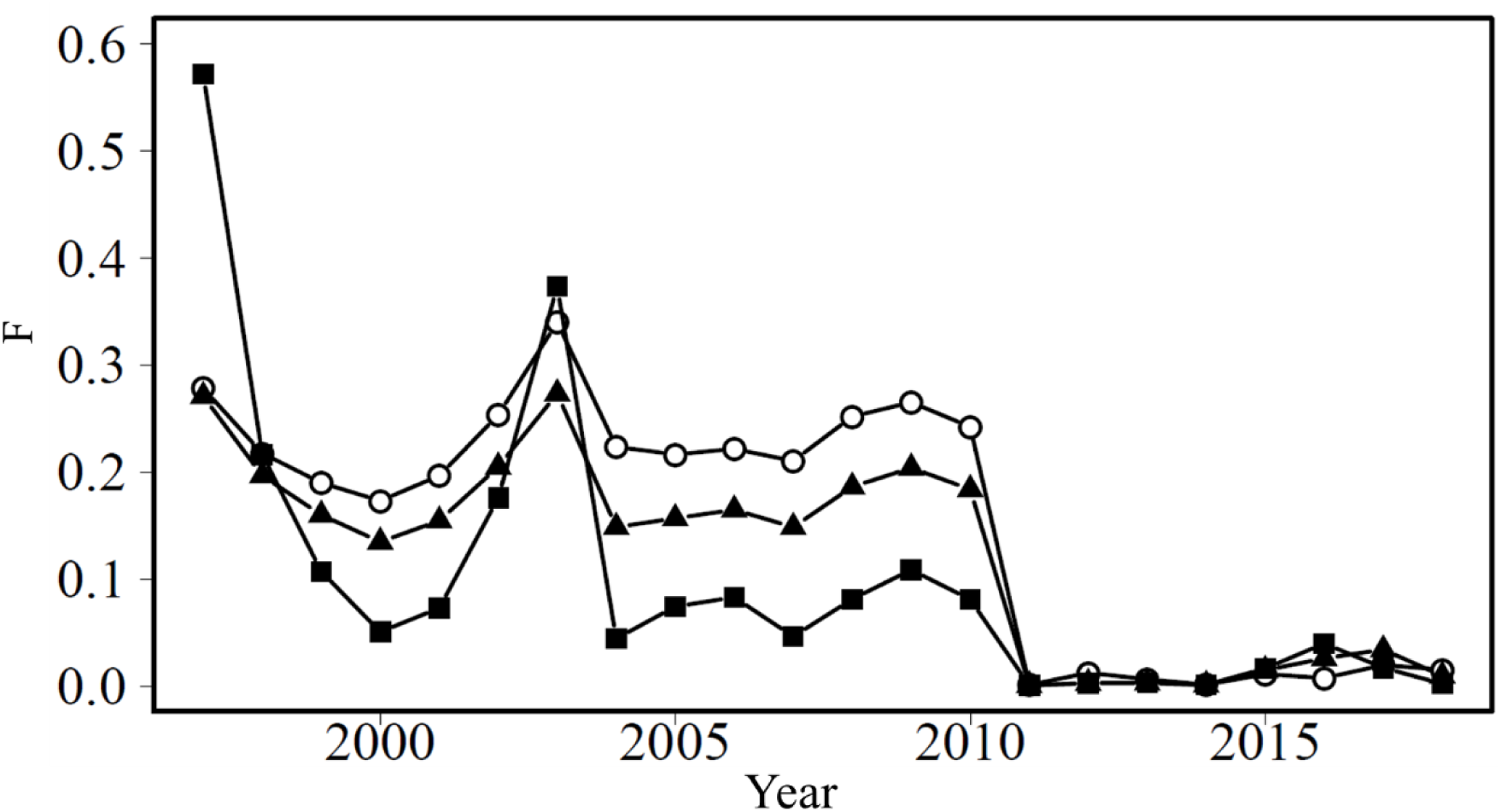
Estimated fishing mortalities of the best model by immature male (black squares), mature male (black triangles), and mature female (white circles).

### Sensitivity analysis of CV

The result showed that the *M_g_*_,*t*_ did not change greatly even if CV was multiplied by 1.5 (Figure 9). The *ρ_past_* and *ρ_future_* changed by this multiplication from 1.2% to –1.8% and from 3.1% to –6.1%, respectively (see also Table 3). This result showed that the estimates and predicted values were robust against the CVs of the number of snow crabs observed by the scientific bottom trawl survey.

**Fig. 9.**
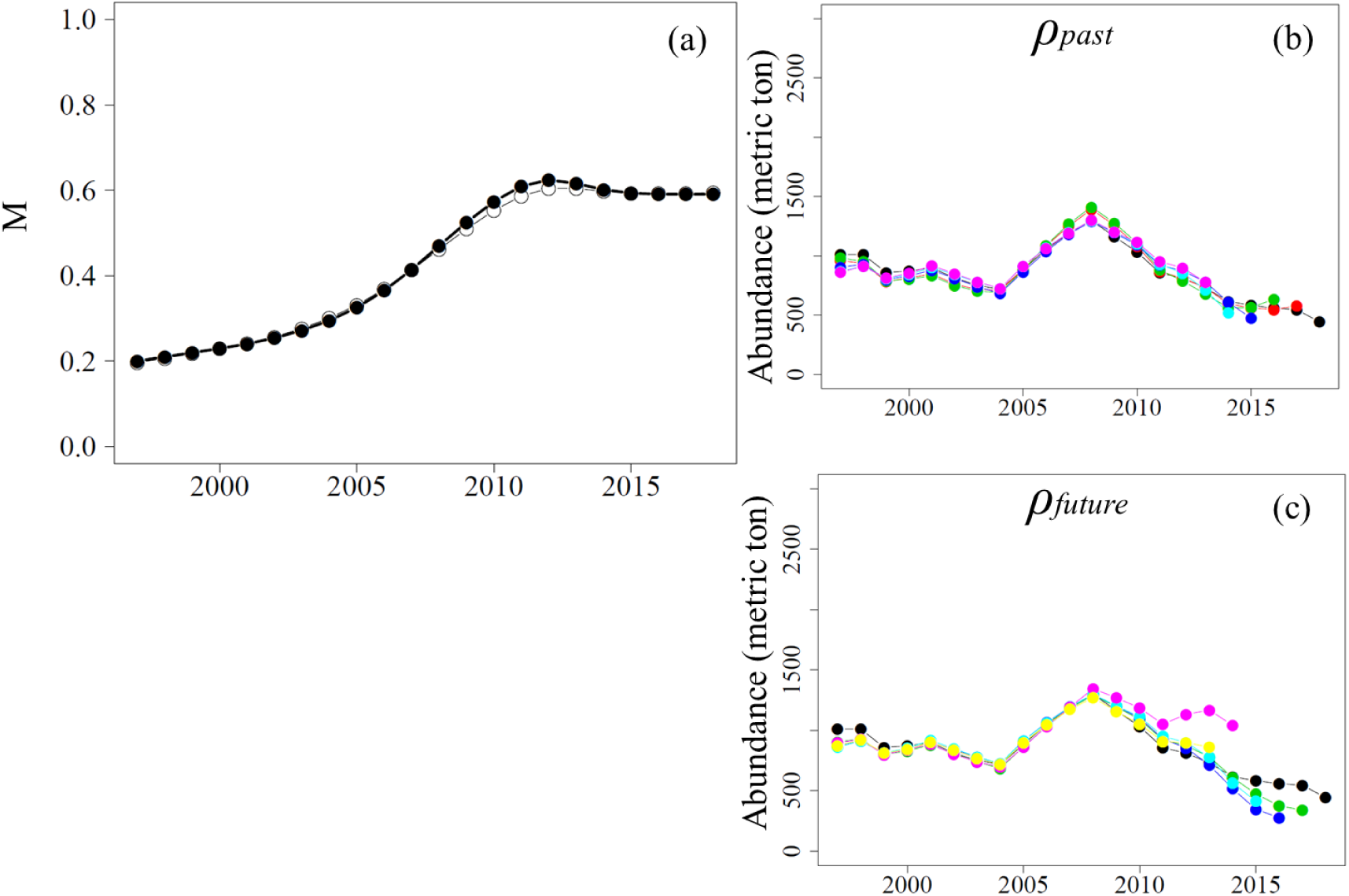
Estimated *M_g,t_* (a) and calculated *ρ_past_* (b) and *ρ_future_* (c) after CV in eq. (32) was multiplied by 1.5. The black and white circles in Fig. 9(a) show values before and after the multiplication, respectively.

### Estimated MSY

The HS model had the minimal AICc because the calculated AICc values were 29.9, 32.21, and 32.0 for the HS, BH, and RI SR models, respectively. The SR relationship estimated by the HS model and estimated *ρ_SR_* are shown in Figure 10a and 10b, respectively. Although the number of instar VIII crabs as recruits decreased since 2015 (Figure 10a), the timing of the decrease was different from that of the abundance (Figure 2). Because the estimated coefficient of autocorrelation was significantly different from zero if the lag was one year (Figure 10b), we used the HS model with autocorrelation in the residuals for future predictions to estimate MSYs based on the three scenarios. Here, the estimated MSY and SSB_MSY_ are shown in Table 5. Although scenario 1 showed that the MSY and SSB_MSY_ were quite high values that had not been experienced in the historical catch (Figure 2) and estimated SSB (Figure 7), both MSY and SSB_MSY_ in scenario 2 were almost zero because the abundance was also almost zero. In scenario 3, the MSY was low compared to the observed historical catch, although the estimated SSBmsy was near to that of the median value from 1997 to 2018 (185.8 gross ton).

**Fig. 10.**
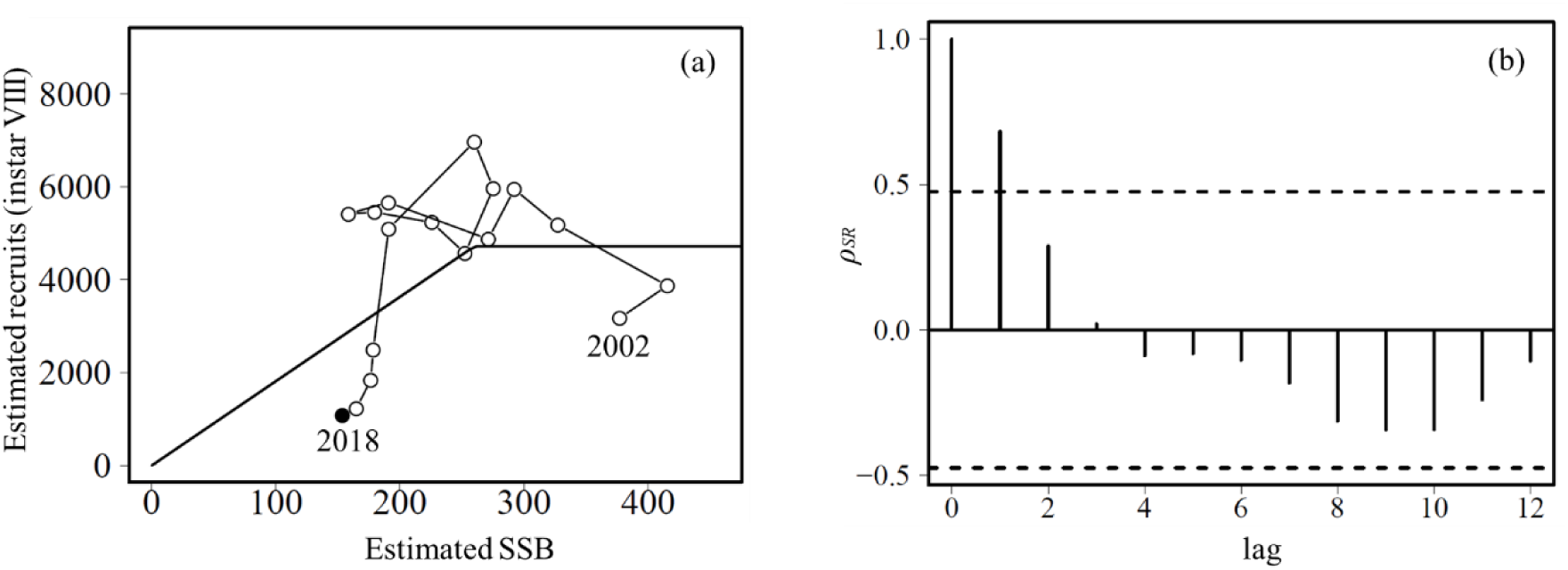
Estimation of the best SR relationship (a) and autocorrelation of their residuals (b) with 95% confidential intervals (break lines).

**Table 5.**
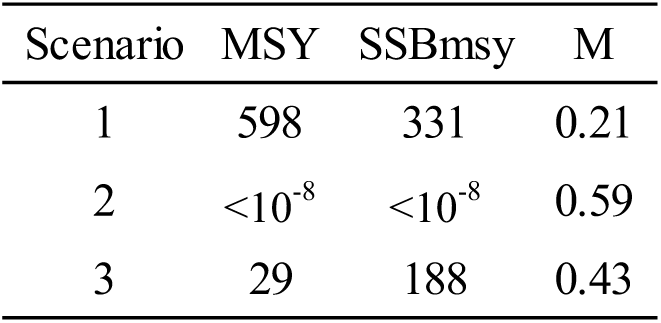
Estimated MSY and SSB_MSY_ for each scenario with their *M* values.

## Discussion

In this study, we showed that the natural mortality coefficient *M* and the terminal molting probability *p* of snow crab in the Tohoku region have increased since 1997. In their stock assessments, it was assumed that *M* = 0.35 for individuals that have not experienced a terminal molt and for those that have experienced a terminal molt within one year, and *M* = 0.2 for those that have undergone a terminal molt two or more years ago (Shibata et al., 2019). In contrast, regardless of the time elapsed since the terminal molt, it has been found in this study that *M* was around 0.59 as of 2018. This means that previous stock assessments overestimated future survival. Indeed, it had been expected that abundance would increase since 2011 because of the rapid decline in total catch (Shibata et al., 2019), although the abundance has maintained a declining trend (Figure 6) despite *F* being nearly zero (Figure 8). Since *F* has been nearly zero and the scientific bottom trawl survey covered the whole habitat of snow crab off Tohoku, the result of this estimation is naturally that *M* and the *p* have increased.

One potential cause for the increase of *M* could be an increased bottom water temperature (BWT). The BWT data were not used in this model since the periods of the surveys were different from those of BWT and we were interested in time-varying parameters thorough the bottom trawl survey period. In contrast, it was suggested that the mean BWT in the Tohoku region was on an upward trend (Figure 11) and a method to draw Figure 11 is shown in Supporting Information 3. An aquarium study indicated that the energy consumed inside the crab exceeds the energy absorbed from outside at a water temperature of 7 °C; therefore it would be energetically impossible for the crab to live persistently in this water temperature range (Foyle et al. 1989). In other words, the increased BWT off Tohoku could be one reason for the increase in *M*. Although the main fishing ground for snow crab has been concentrated in the area from the Miyagi to Ibaraki prefectures (Nemoto 2007), the area of unsuitable environment for survival in the major distribution area of snow crab could have expanded (Figure 11). In contrast, since the trends of BWT and *M* do not completely match, it seems that factors other than the BWT are affecting *M*.

**Fig. 11.**
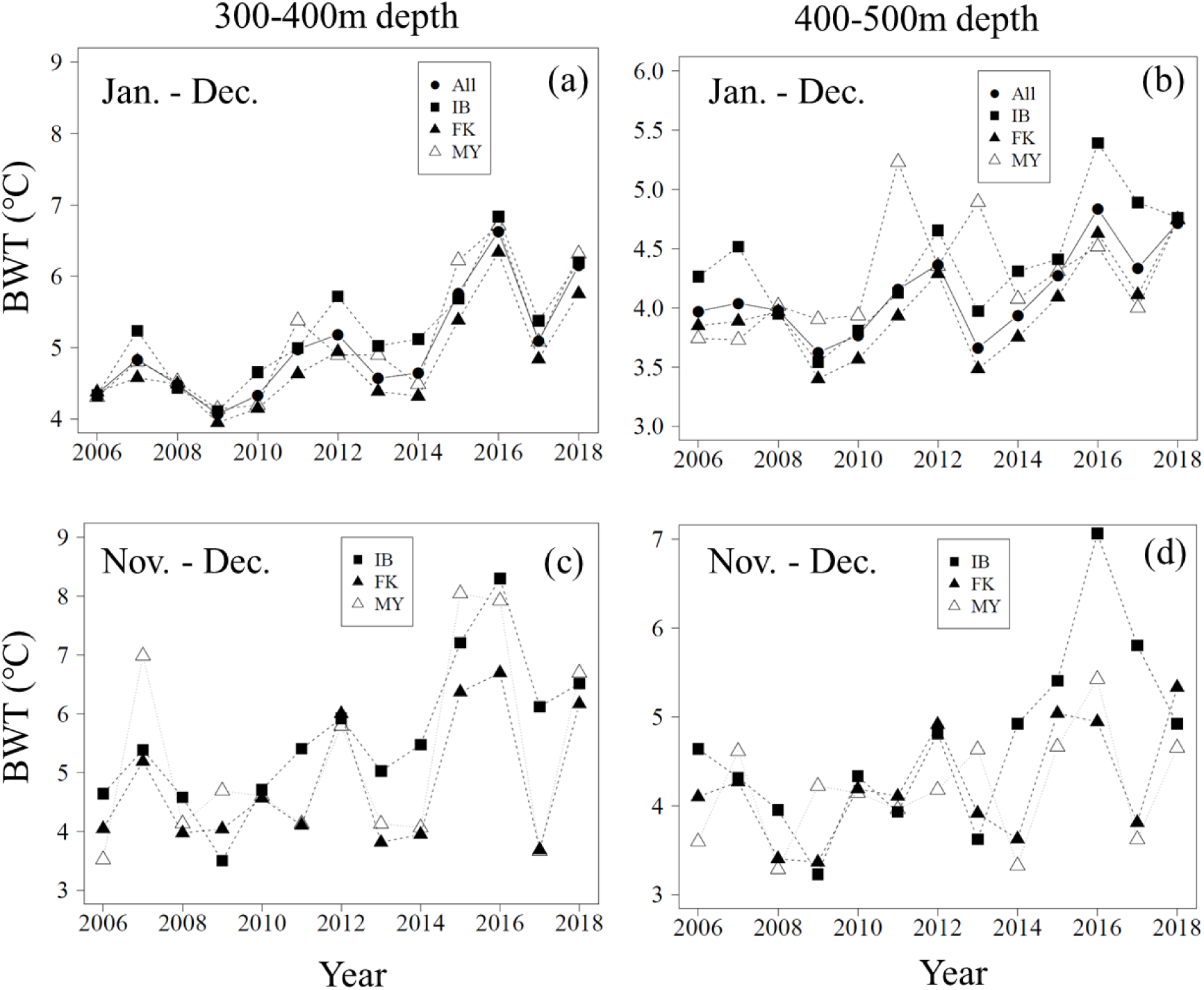
Estimated mean bottom water temperatures (BWT, °C) from January to December (i.e., 12 months) at depths 300–400 m (a) and 400–500 m (b). Mean from November to December (i.e., two months) was also calculated for each depth range (c and d) because the period was the warmest off Ibaraki prefecture (the southern limit of snow crab distribution in the Tohoku region).

Another possible reason for the increase in *M* is predation pressure. It has been known that predation pressure by Atlantic cod (*Gadus morphua*) affects the abundance of snow crab (Chabot et al. 2008). In the Tohoku region, the abundance of Pacific cod (*Gadus macrocephalus*) had increased rapidly since fishing pressure was decreased by the earthquake (Narimatsu et al. 2017) and snow crab have been observed in their stomach contents; therefore, this may have affected the abundance of snow crab (Ito et al. 2014). In contrast, the estimated natural mortality coefficient *M* was the same for all age groups in this study. *M* should differ between small and large snow crabs if the predation pressure by Pacific cod affects the rise in *M*. In fact, it has been reported that snow crabs larger than 65.1 mm rarely appear in the stomach of Atlantic cod (Chabot et al. 2008). The abundance of Pacific cod peaked in 2015 and started to decrease (Narimatsu et al. 2019); however, that of instar VIII snow crab has not turned to an increasing trend, but rather decreased (Figure 10a). This indicates that the predation pressure of Pacific cods is not a main reason for the increase in *M* of all instars.

Not only the natural mortality *M*, but also the terminal molting probability *p*, had increased. It has been reported that the terminal molting probability of snow crab off Miyagi and Fukushima prefectures were higher than that off Ibaraki Prefecture; this might be due to the fact that large individuals were selectively caught under high fishing pressure, resulting in genetically smaller maturity size (Takasaki and Tomiyama 2017). In contrast, this study showed that *p* has been on an upward trend even after 2011, when *F* decreased rapidly because of the earthquake. This therefore suggested that the terminal molting probability would not fluctuate only by fishing mortality. One hypothesis is that recent increases in BWT may affect *p*. Although several studies have reported a positive correlation between water temperature and size-at-terminal molt (Somerton 1981; Alunno-Bruscia and Sainte-Marie 1998; Zheng et al. 2001; Orensanz et al. 2007; Burmeister and Sainte-Marie 2010; Dawe et al. 2012; Yamamoto et al. 2015); however, the Tohoku region has higher water temperatures than any previous studies (Figure 11). It will be necessary to examine the size-at-terminal molt in relatively high-water temperatures through an aquarium experiment.

Our study revealed that the value of MSY apparently changed when the values of *M* and the terminal molting probability changed in three scenarios (Table 5). This suggests that the MSY as an ecosystem service varies greatly with time, and that fishing pressure needs to be reduced to almost zero when *M* and *p* are high. It has been reported that snow crab could have a warm and cold regime for recruitment in the eastern Bering Sea and show drastic stock fluctuation (Szuwalski and Punt 2012, 2013). This study revealed, even in the Tohoku region, the possibility of dynamic stock fluctuations regardless of changes in fishing pressure, because *M* and terminal molting probability varied with time. Although recruits were decreased since 2015 (Figure 10a), this was not the main reason for decreased abundance (Figure 2) because instar VIII as recruits needed at least three more years to be fishable (i.e., instar XI). In other words, the decreased trend in abundance since 2012 was not caused by the decreased recruits since 2015, but by *M* and *p*. Although the catch of snow crab has been limited because bottom trawlers in Fukushima prefecture voluntarily decreased the effort of fishing, the effort should not increase in the future if *M* and *p* maintain high values.

Because the study design allowed for the estimation of *M* and *p*, we could show that ecosystem services can vary with time. Although changes in production due to global warming have been pointed out around the world (Free et al. 2019), it could affect biological parameters, such as *M* and *p*. There could be many species other than snow crab whose biological parameters vary with the environment in the seas around Japan, but there are few examples where the biological parameters have been actually estimated. Japan is currently moving to a new stock management targeting MSY, but it is commonly assumed that the value of *M* has mainly been based on empirically derived equations and does not change over time (Tanaka 1960). For species that are likely to be highly affected by the environment, scientific survey designs to estimate abundance should be prepared to allow the estimation of the parameters and capture of the temporal changes.

One of the features of JASAM is that it estimates parameters for population dynamics using the annual abundance observed independently of fisheries. It is conceivable that the parameters will have other specifications, such as an RW with a multivariate normal distribution, as in the case of *F*, and a step function. It is also possible to change the state model for snow crab to another age-structured model so that it can be applied to species other than snow crab. However, if the estimated abundance does not reflect the entire distribution area of the target species, *M* can be confounded with migration rates to adjacent sea areas. In this case, it may be necessary to assume the rates of migration to the off-site area separately, or to design a survey (e.g., estimation of abundance in the adjacent sea areas) that can estimate the rates of migration.

## Acknowledgment

We thank the crew of the Wakataka-maru, Tanshu-maru, and Hokko-maru for their assistance in obtaining samples. We also thank the staff of Hachinohe Laboratory, Tohoku National Fisheries Research Institute, and Fukushima Prefectural Research Institute of Fisheries Resources for help in preparing the samples. Dr Nishijima gave useful comments for methods to use the TMB. Mr Takahashi, Mr Miharu, Mr Matsumoto, Mr Sato, and Mr Kaneko gave useful comments from the viewpoint of the bottom trawl fishers of Fukushima. This study was funded by the Fisheries Agency of the Ministry of Agriculture, Forestry, and Fisheries of Japan.

## Supporting Information 1

### Correction of CVs using Taylor’s power law

In a swept area method, the mean and standard error are equal when there is a sample at only one station and no samples are obtained at other stations. The situation is simply shown in R code as below:

> x <- c(5, 0, 0, 0, 0, 0)

> mean(x)

[1] 0.8333333

> sd(x)/sqrt((length(x)))

[1] 0.8333333

Although the coefficient of variation (CV) was calculated as one in this case, the value could be higher than expected. We used the CVs corrected by Taylor’s power law (Taylor 1961) as the below regression model:

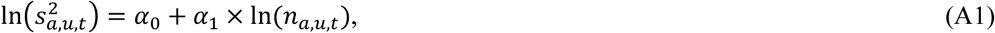

where *n* is the observed number of snow crabs, *s* is their standard error, and the subscripts *a*, *u*, and *t* show the same as in the article. The estimates were 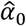 = 0.828 (SE = 0.464), 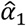 = 1.772 (SE = 0.035), adjusted *R*^2^ = 0.88 and the CVs were corrected using eq. A1 (Fig. A1). There was no change in the best model before and after the correction.

**Figure A1.**
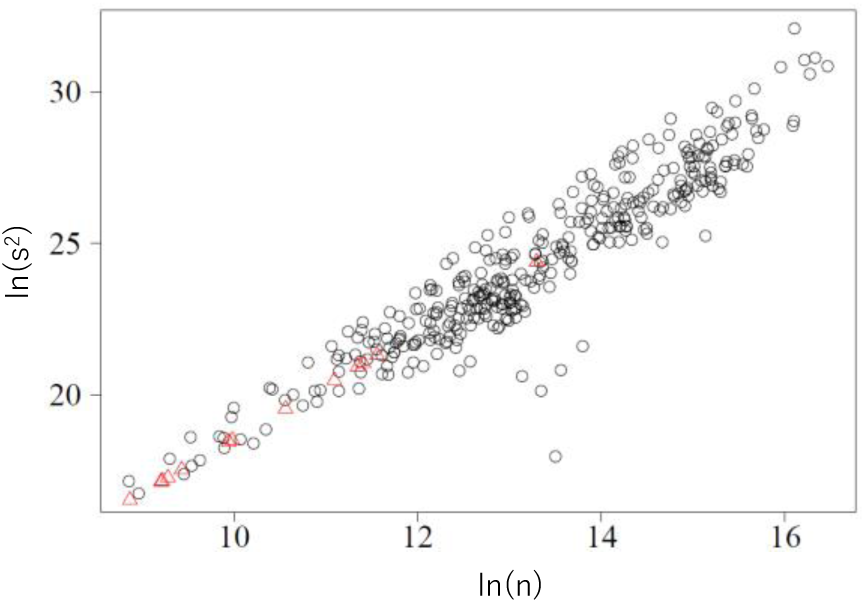
The relationship between ln(n) and ln(s^2^) (black circle). Red triangles show the corrected standard errors using eq. A1.

## Supporting Information 2

### Combinations of age groups

All the 32 combinations of age groups. If the numbers are the same in a combination, those instars (*a=*8*,…,*13) are included in the same age group. *Mg* equals *Ma* in the last combination (i.e., all the instars take a different *M*).

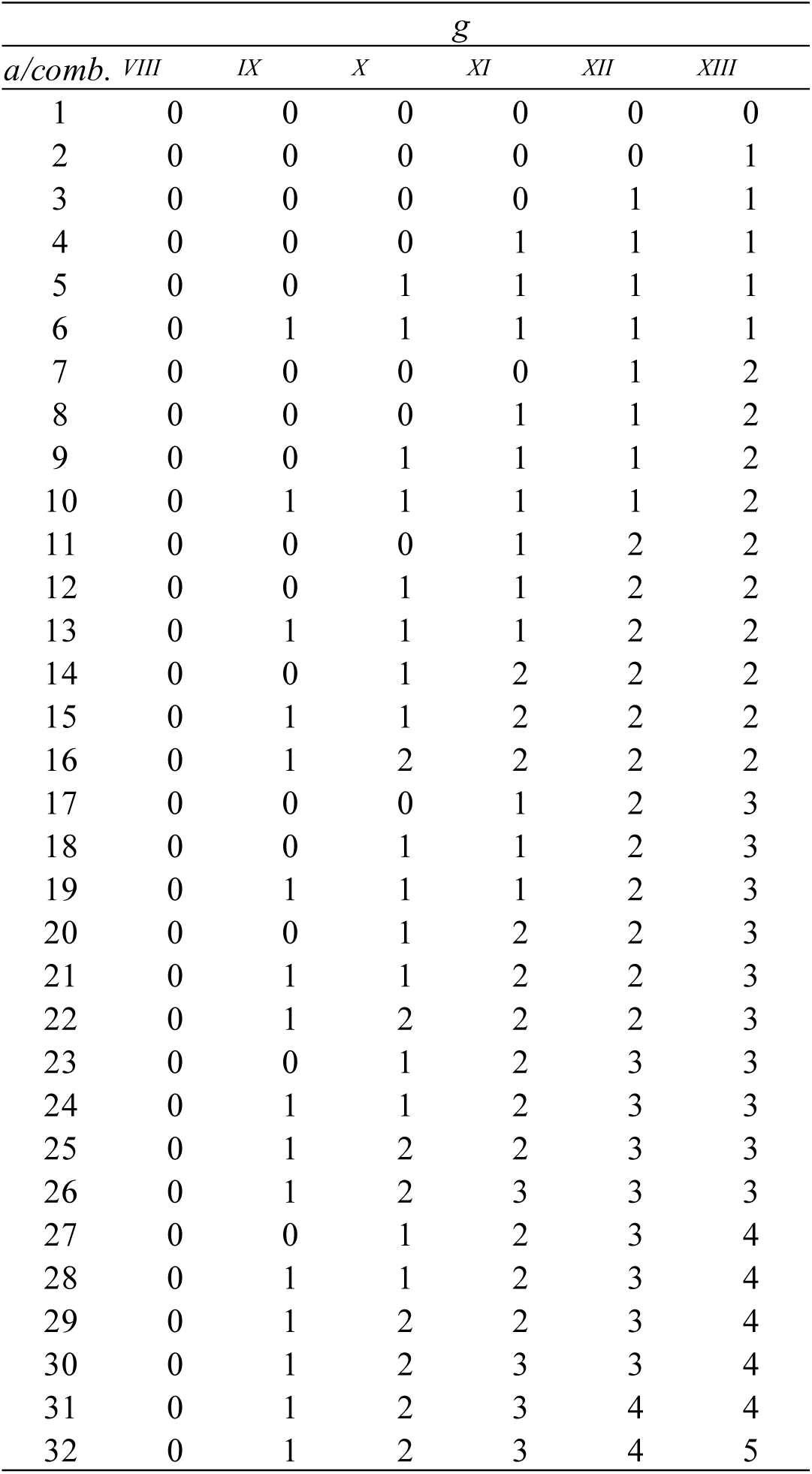

## Supporting Information 3

### Result of fittings for observation

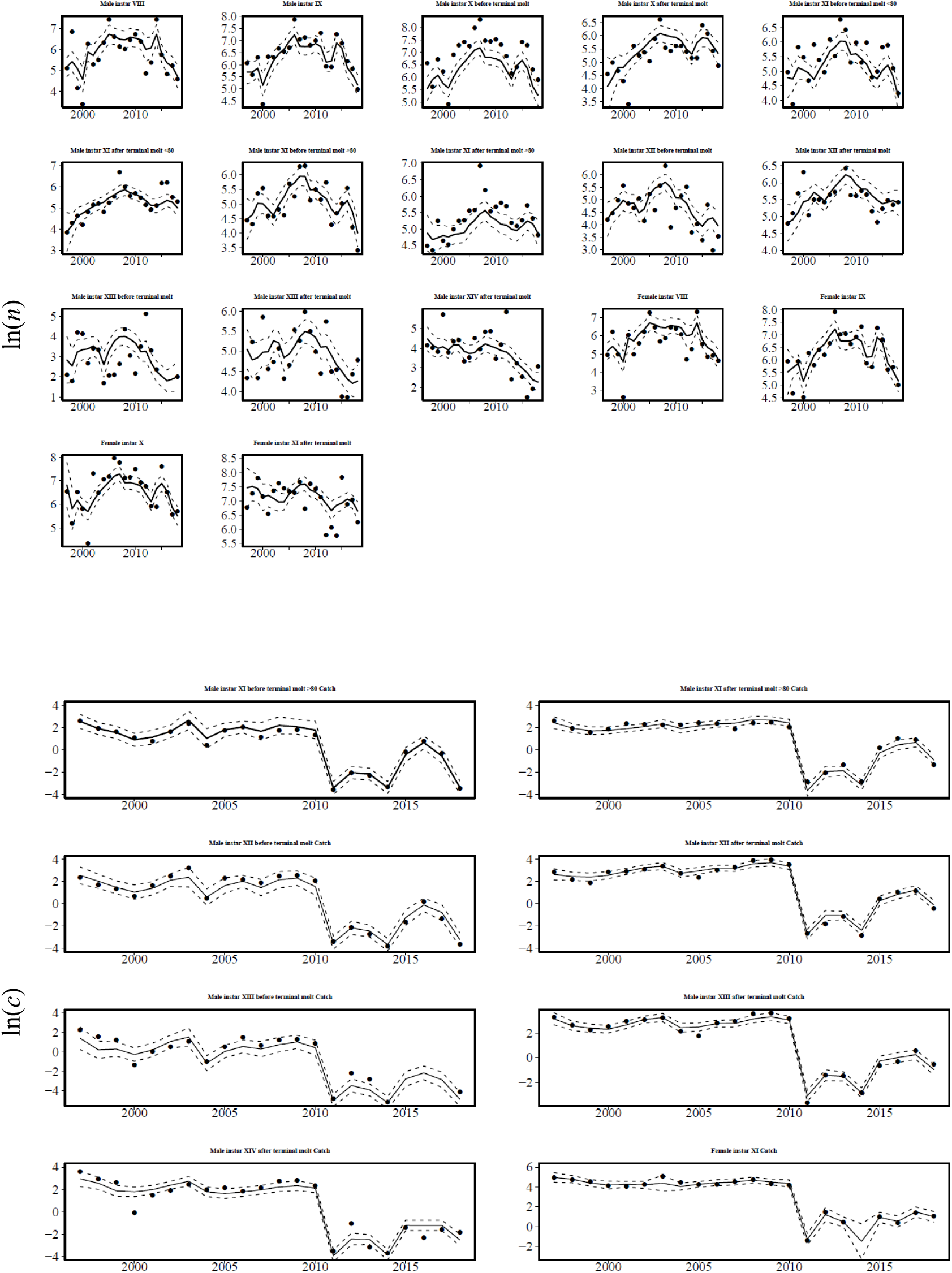

## Supporting Information 4

### Estimated bottom water temperatures

Water temperatures at the maximum observed depth in the Tohoku region were extracted from the conductivity, temperature, and depth (CTD) data obtained by prefectural fisheries experimental stations and Fisheries Research and Education Agency in Japan. Then, we adopted bottom water temperatures using the observed depth of water temperatures as follows: when the depth of the station was ≤ 100 m, the temperatures of which the observed depth was within 10 m from the seabed were adopted. Temperatures of which the observed depth was within 10% of the bottom depth of the seabed were adopted when the depth of the station was > 100 m. The obtained bottom water temperatures were interpolated to monthly gridded data of 5 × 5 min meshes weighed by time, distance, and depth using a flexible Gaussian filter (Shimizu and Ito 1996).

